# The IDR-containing protein PID-2 affects Z granules and is required for piRNA-induced silencing in the embryo

**DOI:** 10.1101/2020.04.14.040584

**Authors:** Maria Placentino, António Miguel de Jesus Domingues, Jan Schreier, Sabrina Dietz, Svenja Hellmann, Bruno F. M. de Albuquerque, Falk Butter, René F. Ketting

## Abstract

In *Caenorhabditis elegans*, the piRNA (21U RNA) pathway is required to establish proper gene regulation and an immortal germline. To achieve this, PRG-1-bound 21U RNAs trigger silencing mechanisms mediated by RNA-dependent RNA polymerase (RdRP)-synthetized 22G RNAs. This silencing can become PRG-1-independent, and heritable over many generations. This state is named RNAe. It is unknown how and when RNAe is established, and how it is maintained. We show that maternally provided 21U RNAs can be sufficient to trigger RNAe in embryos. Additionally, we identify the IDR-containing protein PID-2, as a factor required to establish and maintain RNAe. PID-2 interacts with two novel, partially redundant, eTudor domain proteins, PID-4 and PID-5. Additionally, PID-5 has a domain related to the X-prolyl aminopeptidase protein APP-1, and binds APP-1, implicating N-terminal proteolysis in RNAe. All three proteins are required for germline immortality, localize to perinuclear foci, affect Z granules, and are required for balancing of 22G RNA populations. Overall, our study identifies three new proteins with crucial functions in the *C. elegans* small RNA silencing network.

## Introduction

Germ cells are responsible for transmitting genetic information to the next generation. Therefore, genome stability should be tightly controlled in these cells. The integrity of the genome is constantly threatened by external factors, such as irradiation and mutagenic agents, but also by intrinsic factors resident in the genome, such as transposable elements (TEs). Consequently, organisms have evolved a variety of mechanisms to counteract these threats. Among these, small RNA pathways often play important roles in controlling TE activity. In many animals, TEs are recognized and silenced in the germline by a specific small RNA pathway: the Piwi pathway. Piwi proteins represent a specific subclade of Argonaute proteins that exert their silencing function upon loading with their cognate small RNA, named piRNA (Piwi-interacting RNA), that specifies the target transcript. The Piwi/piRNA pathway is essential in most organisms for TE silencing, but also TE-unrelated effects have been well described (Ghildiyal & Zamore, 2009; Ketting, 2011; Malone & Hannon, 2009; Ozata et al., 2018; Siomi et al., 2011).

The main, and likely only active Piwi protein of *C. elegans* is PRG-1; it binds to piRNAs, that in *C. elegans* are named 21U RNAs, to form a silencing complex. In contrast to other organisms, loss of the PRG-1/21U RNA pathway in *C. elegans* causes the reactivation of only a limited set of transposable elements, for instance Tc3 (Das et al., 2008), and does not cause immediate sterility (Batista et al., 2008; Cox et al., 1998; Das et al., 2008; Wang & Reinke, 2008), even though germ cells are progressively lost over generations (mortal germline phenotype, Mrt) (Simon et al., 2014). The discrepancy between the Piwi-mutant phenotypes observed in *C. elegans* and other animals can be explained by the fact that PRG-1 initiates a silencing response that is executed by a different set of Argonaute proteins – the worm-specific Argonaute proteins (WAGOs) – while in other studied model systems this does not happen. Upon target recognition by PRG-1, an RNA-dependent RNA polymerase (RdRP) is recruited to the targeted transcript, which is used as a template for the synthesis of complementary small RNAs, named 22G RNAs. For this step, the RdRP RRF-1 is required, as well as so-called mutator proteins (Phillips et al., 2014; Phillips et al., 2012; Zhang et al., 2011). The 22G RNAs, characterized by the 5’ triphosphate group resulting from the RdRP-driven synthesis, are loaded onto WAGO proteins, such as HRDE-1 and WAGO-1 (Ashe et al., 2012; Buckley et al., 2012; Gu et al., 2009; Shirayama et al., 2012), that reinforce the silencing started by PRG-1. Occasionally, in a seemingly stochastic and poorly understood manner, this silencing can become independent of PRG-1 itself, and self-sustainable. This form of silencing is extremely stable and can be maintained across many generations in absence of PRG-1. It is characterized by the deposition of heterochromatic marks at the targeted locus, depends on HRDE-1 and mutator activity, and it is known as RNAe (RNA-induced epigenetic gene silencing) (Ashe et al., 2012; Luteijn et al., 2012; Shirayama et al., 2012). RNAe can thus explain why transposons remain silenced in absence of PRG-1. Indeed, in *prg-1;hrde-1* double mutants, lacking both 21U RNAs and RNAe, the activity of Tc1 transposons increases to levels comparable to mutator mutants, indicating that HRDE-1 activity is sufficient to maintain Tc1 silencing in *prg-1* mutants (de Albuquerque et al., 2015).

PRG-1/21U RNA complexes can recognize a target transcript via imperfect base-pair complementarity, allowing up to 4 mismatches (Bagijn et al., 2012; Lee et al., 2012). As a consequence of this mismatch tolerance, PRG-1 is potentially able to recognize and silence many different sequences, including endogenous genes (Bagijn et al., 2012; Gu et al., 2012). Another small RNA pathway, guided by 22G RNAs bound to the Argonaute protein CSR-1, has been implicated in counteracting such PRG-1-mediated silencing of genes that should be expressed (Claycomb et al., 2009; Conine et al., 2013; Gu et al., 2009; Lee et al., 2012; Seth et al., 2013; Shen et al., 2018; Shirayama et al., 2012; Wedeles et al., 2013). CSR-1-bound 22G RNAs are made by the RdRP EGO-1 in a mostly mutator-independent manner (Claycomb et al., 2009; Gu et al., 2009). Interestingly, an opposite scenario has also been described: PRG-1 has been shown to direct mutator activity to non-CSR-1 targets in embryos that set up a 22G RNA silencing response *de novo* (de Albuquerque et al., 2015; Phillips et al., 2015). These seemingly contradictory findings – CSR-1 counteracting inappropriate PRG-1 targeting versus PRG-1 directing mutator activity away from CSR-1 targets – may be explained by considering that two different developmental stages have been analysed to arrive at the proposed models. The protective role of CSR-1 has been seen in the adult germline (Claycomb et al., 2009; Conine et al., 2013; Gu et al., 2009; Lee et al., 2012; Seth et al., 2013; Shen et al., 2018; Shirayama et al., 2012; Wedeles et al., 2013), whereas the protective role of PRG-1 likely operates in embryos (de Albuquerque et al., 2015; Phillips et al., 2015). Possibly, PRG-1 has different modes of actions at these two developmental stages. Another result that points in this direction comes from studies on HENN-1, the enzyme that 2’-O-methylates 21U RNAs. In adults, 21U RNA levels are not affected by loss of HENN-1 (Kamminga et al., 2012), while in embryos 21U RNAs are reduced in *henn-1* mutants (Billi et al., 2012; Montgomery et al., 2012). Given that 2’-O-methylation has been shown to stabilize small RNAs, in particular when they base-pair extensively to their targets (Ameres et al., 2010), it is feasible that PRG-1 recognizes targets with near-perfect complementarity to its cognate 21U RNA only in the embryo, and employs more relaxed 21U RNA-targeting in the adult germline. Indeed, maternally provided PRG-1 protein is required to establish PRG-1-driven silencing of a 21U RNA sensor transgene that has perfect 21U RNA homology, suggesting that this silencing is set up during early development, and not in the adult germline (de Albuquerque et al., 2014). Whether the maternal contribution of PRG-1 is sufficient to induce silencing has not been tested thus far.

A third small RNA pathway is driven by so-called 26G RNAs (Billi et al., 2014; Conine et al., 2010; Han et al., 2009; Yigit et al., 2006). These are made by the RdRP enzyme RRF-3, which acts in a large protein complex containing well conserved proteins such as Dicer, GTSF-1 and ERI-1 (Almeida et al., 2018; Billi et al., 2014; Duchaine et al., 2006; Kennedy et al., 2004; Thivierge et al., 2012). These 26G RNAs can be bound by the Argonaute protein ERGO-1, or by two closely related paralogs, the Argonaute proteins ALG-3 and ALG-4 (ALG-3/-4). ERGO-1 mostly targets transcripts in the female germline and the early embryo, and is required to load the somatic, nuclear Argonaute protein NRDE-3 with 22G RNAs (Almeida et al., 2019a; Billi et al., 2014; Gent et al., 2010; Han et al., 2009; Vasale et al., 2010). The 26G RNAs bound by ERGO-1 require HENN-1-mediated 2’O-methylation in both the adult germline as well as in the embryo (Billi et al., 2012; Montgomery et al., 2012; Kamminga et al., 2012). ALG-3/-4-bound 26G RNAs are not modified by HENN-1 (Billi et al., 2012; Montgomery et al., 2012; Kamminga et al., 2012) and are specifically expressed in the male gonad (Conine et al., 2010; Conine et al., 2013; Han et al., 2009).

Many of the above-mentioned proteins are found in phase-separated structures, often referred to as granules or foci. Mutator proteins that make 22G RNAs are found in so-called mutator foci, whose formations is driven by MUT-16, a protein with many intrinsically disordered regions (IDRs) (Phillips et al., 2012; Uebel et al., 2018). The RdRP EGO-1, as well as the Argonaute proteins CSR-1, PRG-1 and a number of others, are found in P granules (Batista et al., 2008; Claycomb et al., 2009; Updike & Strome, 2010; Wang & Reinke, 2008), characterized by IDR proteins such as PGL-1 (Kawasaki et al., 1998) and DEPS-1 (Spike et al., 2008), which are also required for P granule formation. Finally, Z granules are marked by the conserved helicase ZNFX-1 and the Argonaute protein WAGO-4 (Ishidate et al., 2018; Wan et al., 2018). Z granules are related to the inheritance of small RNA-driven responses via the oocyte (Ishidate et al., 2018; Wan et al., 2018) and are typically found adjacent to P granules. However, in primordial blastomeres, Z and P granules appear to be merged (Wan et al., 2018). For Z granules, no IDR protein that may drive their formation has been identified yet. The function of ZNFX-1 is also not clear, but it has been demonstrated that it interacts with the RdRP EGO-1, and that it is required to maintain the production of 22G RNAs from the complete length of the targeted transcript (Ishidate et al., 2018). In absence of ZNFX-1, relatively more 22G RNAs are found to originate from the 5’ part of the RdRP substrate, suggesting that ZNFX-1 may have a role in maintaining or relocating the RdRP activity to the 3’ end of the substrate. Despite the fact that material exchange between these three types of structures (P, Z granules and mutator foci) seems obvious, how this may happen is currently unknown.

Here, we describe the characterization of a novel gene, *pid-2*, which we identified from our published “piRNA-induced silencing <u>d</u>efective” (<u>Pid</u>) screen (de Albuquerque et al., 2014). Our analyses show that PID-2 is essential for initiation of silencing by maternally provided PRG-1 activity. However, PID-2 is also required for efficient maintenance of RNAe, and shows defects in many different small RNA populations, including 26G RNAs, indicating that PID-2 does not specifically act together with PRG-1. Interestingly, we noticed a drop of 22G RNA coverage specifically at the 5’ end of RRF-1 substrates, suggesting that PID-2 may be involved in stimulating RdRP activity, or processivity. At the sub-cellular level, PID-2 is found in granules right next to P granules, and absence of PID-2 affects size and number of Z granules, suggesting that PID-2 itself may also be in Z granules. We also identify two PID-2-interacting proteins, PID-4 and PID-5, with an extended Tudor (eTudor) domain. In addition, PID-5 has a domain that closely resembles the catalytic domain of the X-prolyl aminopeptidase protein APP-1. Loss of both PID-4 and PID-5 phenocopies *pid-2* mutants in many aspects, including the effect on Z granules. However, at steady-state, both PID-4 and PID-5 are themselves mostly detected in P granules. We hypothesize that the identified PID-2/-4/-5 proteins have a role in controlling RdRP activity, and do so by affecting protein and/or RNA exchange between different phase-separated granules.

## Results

### PID-2 is an IDR-containing protein required for 21U RNA-driven silencing

In order to identify novel factors of the 21U RNA machinery, we previously performed and published a forward mutagenesis screen in which we identified several mutants that are defective for 21U RNA-driven silencing (piRNA-induced silencing defective: Pid). As a readout, we used the de-silencing of a fluorescent 21U RNA target (21U sensor), which was not yet in an RNAe state (Bagijn et al., 2012; de Albuquerque et al., 2014). Among the mutants isolated from the screen, we focussed our attention on one strain, characterized by a point mutation (tgg → tga) causing a premature stop codon (W122X) in the gene Y48G1C.1 (Figure 1A). The protein encoded by this gene contains disordered N- and C-terminal regions, while the rest of the protein appears to be more structured (Figure 1A), even though no predicted domains were detected. Animals homozygous for the *xf23* allele cannot fully silence the 21U sensor, even though its expression levels are lower than in mutator mutants, as judged by imaging (Figure 1B). We then crossed a 21U sensor, again in a non-RNAe state, into a strain with a publicly available deletion allele of Y48G1C.1, *tm1614* (Figure 1A) (C. elegans Deletion Mutant Consortium, 2012). This de-silenced the 21U sensor to a similar extent (Figure 1B). We also generated single-copy transgenes expressing a 3xFLAG- or eGFP-tagged version of Y48G1C.1, driven by its endogenous promoter and 3’ UTR in order to rescue the 21U sensor reactivation phenotype (Figure S1A). Both C-terminally eGFP-tagged (*xfIs145*) and N-terminally 3xFLAG-tagged (*xfIs146*) Y48G1C.1 could partially rescue the silencing defect of *xf23* (Figure S1B, C, D). To quantify the de-silencing induced by *tm1614* and *xf23* we used RT-qPCR, revealing very similar activation of the 21U sensor by both alleles (Figure S1E). We conclude that the mutation in Y48G1C.1 is responsible for the *xf23* phenotype and named the gene *pid-2*.

**Figure 1.**
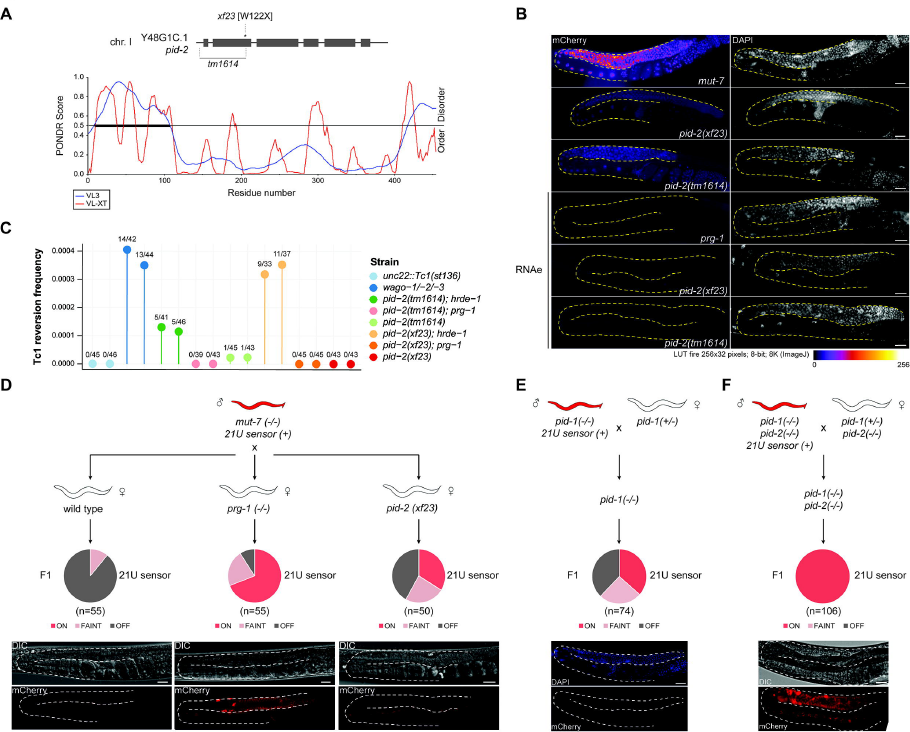
PID-2 is a novel factor required for establishing *de novo* silencing mediated by maternally provided small RNAs. **A)** Schematic representation of the Y48G1C.1 / *pid-2* gene and its mutant alleles (*xf23* and *tm1614*). The line plot displays the predicted disorder of the Y48G1C.1 / PID-2 protein, as obtained from PONDR, using the algorithms VL3 and VL-XT. **B)** Expression of the 21U sensor (left) and DAPI staining (right) of gonad arms in the indicated genetic backgrounds. Gonads are outlined by a dashed line. The mCherry signal is represented in pseudo-colours [LUT fire (ImageJ)] to reflect differences in the intensity of the signal. Scale bar: 25 μm. **C)** Tc1 reversion assay in different genetic backgrounds. All the strain tested carry the *unc-22::Tc1(st136)* allele. Negative control = *unc-22::Tc1(st136)* in a wild type background; positive control = *wago-1/-2/-3*. In each experiment, 50 plates per strain have been initially tested. Two independent experiments are represented. Above each data point, the number of plates with revertants are indicated. The amount of worms has been estimated to be 1000 worms/plate. **D)** Crossing scheme to address the re-initiation of silencing of the 21U sensor. A *mut-7* mutant male expressing the 21U sensor is crossed with either a wild type (left), *prg-1* (middle) or *pid-2* (right) mutant hermaphrodite. Their F1 offspring was scored for expression of the 21U sensor by microscopy. Three states of expression were scored, and represented in a pie chart. The three expression states are exemplified by representative images at the bottom: silenced (left), faintly expressed (right) or expressed (middle). DIC images are shown above the fluorescence panels. Gonads are outlined by a dashed line. Scale bar: 25 µm. **E)** Crossing scheme to address if maternal 21U RNAs are sufficient to re-initiate the silencing of the 21U sensor. A *pid-1* mutant male expressing the 21U sensor is crossed with a hermaphrodite, heterozygous for the same mutation. All their F1 offspring inherit a pool of 21U RNAs from the hermaphrodite, but in 50% of the F1, which is *pid-1* homozygous mutant, no zygotic PID-1 is present, hence no zygotic 21U RNAs can be made. The silencing or expression of the 21U sensor in the *pid-1* homozygous mutant F1 has been scored by microscopy, and depicted in a pie chart. At the bottom, a representative image of an animal carrying a silenced 21U sensor (lower: mCherry signal; upper: DAPI staining) in *pid-1* mutant offspring. Gonads are outlined by a dashed line. Scale bar: 25 μm. **F)** Crossing scheme to test the role of PID-2 in re-initiating the silencing of the 21U sensor, mediated by maternally provided 21U RNAs only. The expression of the 21U sensor in the F1 has been scored by microscopy, and depicted in a pie chart. DIC and fluorescence image of a representative animal are shown at the bottom. Gonads are outlined by a dashed line. Scale bar: 25 μm.

We also tested whether the *xf23* and *tm1614* alleles reactivate a 21U sensor that is in a state of RNAe, *i.e.* whose silencing had become PRG-1-independent. Our initial experiments showed that neither the *xf23* nor the *tm1614* allele were able to do so (Figure 1B). Nonetheless, an RT-qPCR experiment performed at a later time point showed that the 21U sensor was actually expressed in one of the *pid-2* mutant strains that originally received the 21U sensor in an RNAe state (Figure S1E). This primed us to investigate this 21U sensor (RNAe) reactivation more thoroughly. We generated clonal populations from *prg-1*, *pid-2* and *prg-1;pid-2* mutant individuals, all carrying the 21U sensor in an RNAe state and followed them over time, both at 20 ^o^C and at 25 ^o^C. After several generations, we noticed that the 21U sensor was indeed reactivated in some of the *pid-2* mutants at 20 ^o^C, and even more frequently at 25 ^o^C, whereas we never observed such reactivation in *prg-1* mutants (Figure S1F). Even though from these data we cannot draw quantitative conclusions, they do show that loss of PID-2 cripples the efficiency of RNAe maintenance. Strikingly, we did not observe any reactivation of the 21U sensor in a *prg-1;pid-2* double mutant (Figure S1F). We can currently not explain this effect, but it is possible that PRG-1 and the RNAe machinery compete for shared resources, involving PID-2: in a *prg-1;pid-2* double mutant background, RNAe may become “stronger” because PRG-1 does not draw on these resources. We note that Tc1 re-activation shows similar characteristics (see below). Overall, we conclude that PID-2 is involved in both the initiation, through PRG-1, as well as the maintenance of RNAe on a 21U RNA-silenced transgene, but also that some silencing can still occur in absence of PID-2.

### PID-2 acts together with HRDE-1 to silence Tc1 transposition

Previously, we showed that loss of PRG-1 can trigger Tc1 activity in *hrde-1* mutants (de Albuquerque et al., 2015). This indicates that both PRG-1 and HRDE-1 are required to silence Tc1 and that each of them alone is able to suppress Tc1 activity. Hence, we tested whether *pid-2* mutation impairs Tc1 silencing, by itself or in combination with *prg-1* or *hrde-1* mutation. We observed Tc1 reactivation in *pid-2;hrde-1* double mutants, but not in *prg-1;pid-2* double mutants (Figure 1C). In fact, no reversions at all were seen on any of the plates with *prg-1;pid-2* double mutants, in contrast to *pid-2* single mutants, in line with the above-described lack of RNAe loss in *prg-1;pid-2* double mutants. Our interpretation is that the crippled PRG-1-driven silencing of *pid-2* mutants is not sufficient to control Tc1 activity in absence of HRDE-1, whereas the HRDE-1-driven silencing of Tc1 in *pid-2* mutants is still sufficient, and possibly even more robust, in absence of PRG-1.

### Maternally inherited 21U RNAs act via PID-2 to establish *de novo* silencing

We showed before that maternally provided PRG-1 activity is required to initiate silencing of a 21U sensor transgene (de Albuquerque et al., 2014). We repeated this experiment and also included *pid-2* mutants. This revealed that, like PRG-1, PID-2 displays a maternal effect: a significant fraction of the offspring of *pid-2* mutant mothers could not induce silencing on a 21U sensor that was brought in via the sperm, despite the fact that this offspring did carry a wild-type copy of *pid-2* (Figure 1D).

We next wanted to test whether maternal contribution of 21U RNAs could be sufficient for *de novo* target silencing. To do this, we made use of the *pid-1* mutation, which disrupts 21U RNA production (de Albuquerque et al., 2014). We crossed *pid-1* mutant males that express the 21U sensor, with *pid-1* heterozygous hermaphrodites. Offspring from this cross that was again *pid-1* homozygous mutant, could not produce 21U RNAs themselves, but did receive maternal PRG-1 and 21U RNAs. We observed that a large fraction of the *pid-1* mutant offspring (~40%) was able to silence the 21U RNA sensor *de novo*, indicating that the maternal 21U RNAs were sufficient for silencing (Figure 1E). Interestingly, the silencing that was thus established was transmitted stably for many generations. We sequenced small RNAs from strains isolated from these crosses, in which the 21U sensor was either silenced or still active. This revealed absence of 22G RNAs targeting the 21U sensor in non-silenced strains, and a typical RNAe-like 22G RNA pattern in the silenced strains (Figure S1G). The silenced state induced by maternal 21U RNAs was also found to be sensitive to loss of HRDE-1 (Figure S1H). We conclude that maternal 21U RNAs can be sufficient to induce RNAe.

We then introduced a *pid-2* mutation in this setup. We observed that none of the *pid-1* mutant offspring was able to silence the 21U sensor without PID-2, and all displayed strong expression (Figure 1F). Interestingly, all F1 from this cross that was *pid-1;pid-2* homozygous, but not those that were *pid-1/+;pid-2*, showed germline defects. In particular, these animals were feminized, as evidenced by the characteristically arrayed oocytes lined up before the spermatheca (Figure 1F and S1I) and the fact that we could rescue their sterility by mating to a wild-type male. This was surprising, as we were able to maintain a *pid-1;pid-2* homozygous mutant strain. However, upon careful investigation of this strain, we did observe feminized individuals (3/30 animals). In addition, a pseudo-male animal (1/30 animals) was detected (Figure S1I). It has been shown that a specific 21U RNA acts in the sex determination pathway, thereby playing a role in proper gonad development (Tang et al., 2018). While this can be related to our observation, it currently remains unclear why the feminization phenotype was so much more prominent in our crosses than in the established double mutant strain. We conclude that PID-2 is essential for the silencing activity of maternally contributed 21U RNAs, and that absence of PID-2 can reveal a strong feminization phenotype in *pid-1* mutants.

### Loss of PID-2 causes a reduction of 21U sensor-derived 22G RNAs

We next performed small RNA sequencing on gravid adults to uncover defects in small RNA populations caused by the absence of PID-2, which could explain the 21U sensor reactivation in *pid-2* mutants. We sequenced at least triplicates of each strain. First, we checked 21U RNA levels, but found these to be virtually unchanged (Figure S2A). Hence, the 21U sensor silencing defect was not due to loss of 21U RNAs. We then checked 22G RNAs that are derived from the 21U sensor transgene. As controls, we sequenced wild-type animals carrying a silenced sensor (Figure 2: panel I), *prg-1* mutant strains in which the sensor was either expressed or not (RNAe) (Figure 2: panels II, III), and *hrde-1* and *mut-7* mutants, in which the RNAe status was disrupted (Figure 2: panels IV, V). In wild-type animals two populations of 22G RNAs could be seen: one that is close to the indicated 21U RNA recognition site, and one that spreads along the mCherry coding region. The one close to the 21U RNA binding site has been named secondary 22G RNAs (Sapetschnig et al., 2015). They are likely triggered directly by PRG-1, as this population is gone in *prg-1* mutants in which the sensor is active (Figure 2: panel II), but are much less affected by loss of HRDE-1 (Figure 2: panel IV). The pool produced along the mCherry coding sequence has been dubbed tertiary 22G RNAs (Sapetschnig et al., 2015), and was found to be dependent on HRDE-1. Loss of MUT-7 strongly affected both secondary and tertiary 22G RNA pools (Figure 2: panel V). For reasons we do not understand, the *his-58* region was not covered by 22G RNAs.

**Figure 2.**
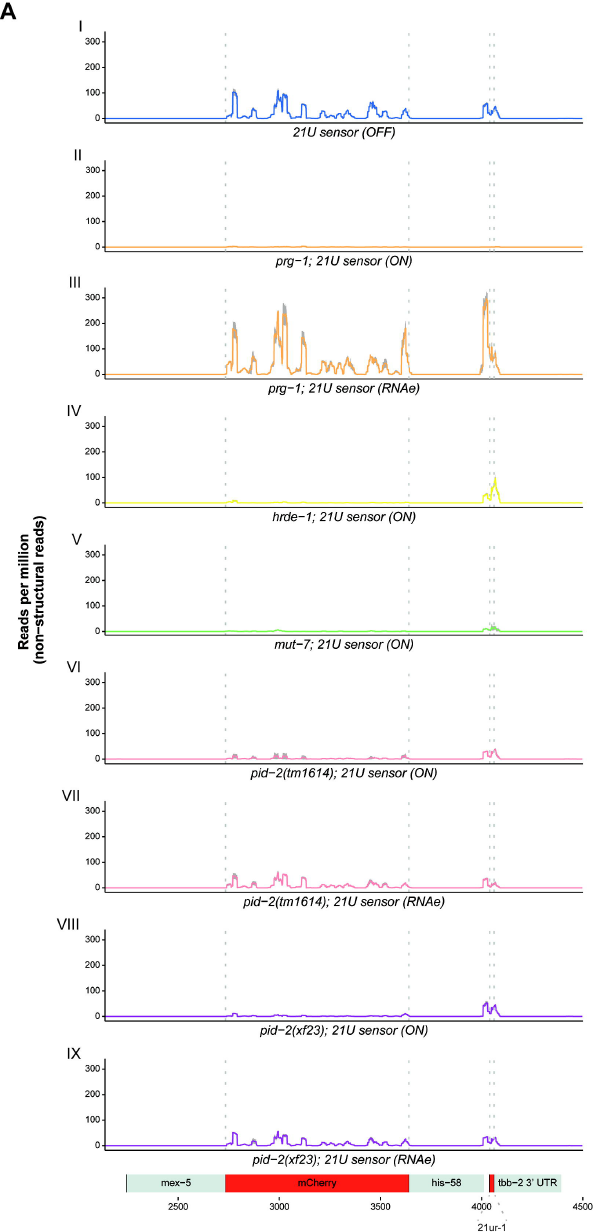
Small RNA sequencing of 22G RNAs mapping to the 21U sensor in *pid-2* mutants. The 22G RNAs mapping to the 21U sensor transgene were identified from total 22G RNA sequencing data, and read density was plotted over the transgene, which is schematically depicted at the bottom. We typically do not detect reads mapping to the *his-58* part of the transgene. The aggregated results of three replicates is shown for each indicated genotype in the different panels (I-IX). The shading, in grey, represents the standard deviation of the read density from the three replicates. ON means that the sensor was expressed. OFF means that the sensor was not expressed, or was crossed into *pid-2* mutant backgrounds in an RNAe state (*i.e.* its silencing was PRG-1-independent).

We then analysed the effect of *pid-2(xf23)* and *pid-2(tm1614)* on these 22G RNA populations. For this, we crossed the 21U sensor into *pid-2* mutants, either from a *mut-7* mutant background in which it was expressed, or from a *prg-1* mutant background in which it was under control of RNAe (Figure 2: panels VI-IX). Both alleles basically produced the same results. First, the secondary 22G RNAs were reduced compared to wild-type and *hrde-1* mutants. This was true whether the transgene originally was under RNAe or not (Figure 2: panels VI-IX; S2B). However, consistently fewer secondary 22G RNAs were detected when the sensor originally was under RNAe, even though this effect was small (Figure 2: panels VII, IX; S2B). This may reflect the fact that fewer transcripts will be available for PRG-1 to recognize when the sensor is silenced by RNAe. Reduced secondary 22G RNA coverage was also found on endogenous PRG-1 target sites (Figure S2C, D). These results show that the direct 22G RNA response to PRG-1 is impaired, but not absent in *pid-2* mutants. Second, tertiary 22G RNAs were almost completely lost when the 21U sensor was introduced in an active state (Figure 2, panel VI, VIII; S2B). In contrast, tertiary 22G RNAs were reduced, but still clearly detectable when the 21U sensor was introduced in an RNAe state (Figure 2, panel VII, IX; S2B). Altogether, we conclude that lack of PID-2 dampens the overall propagation of 22G RNA populations, both secondary as well as tertiary, on the 21U RNA sensor, but does not eliminate it. PID-2 is critically required, though, to initiate an RNAe-related 22G RNA population *de novo*, consistent with its requirement for maternal 21U RNA-induced silencing.

### PID-2 affects endogenous 22G and 26G RNA populations

We also checked the effect of PID-2 on other classes of endogenous small RNAs. As expected, miRNAs were unaffected in *pid-2* mutants (Figure S2A). Interestingly, the strongest effect we observed on total pools of small RNA types was on 26G RNAs (Figure S2A). Consistent with that, also 22G RNAs produced from ERGO-1 targets were reduced in *pid-2* mutants compared to wild type (Figure S2C). The effect on overall bulk 22G RNA levels was only minor (Figure S2A). Also when we split the 22G RNAs into previously defined sub-categories (Almeida et al., 2019a; Bagijn et al., 2012; Conine et al., 2013; Gu et al., 2009; Phillips et al., 2014; Zhou et al., 2014), we only identified relatively small differences in all pathways analysed (Figure S2C).

These bulk analyses are blind to potentially strong effects on individual genes. We therefore performed a differential targeting analysis to identify potential individual genes that either gained or lost 22G RNAs in *pid-2* mutants. This revealed that many genes were consistently either up- or down-regulated in *pid-2* mutants (Figure 3A). Specifically, with a cut-off at 2-fold change and adjusted p-value < 0.05, we detected 1499 genes that lost 22G RNAs and 1482 genes that gained 22G RNAs (Figure 3A; Table S1). We asked if these two sets of genes overlapped significantly with gene sets defined as mutator, CSR-1, ALG-3/-4 or ERGO-1 targets (Figure 3B; S3A-D), and found that the PID-2-responsive genes were enriched for mutator targets. This is consistent with the fact that many mutator targets lost 22G RNAs in *prg-1* mutants (Figure S2C). We can currently not explain why some genes gained and other genes lost 22G RNAs, but this effect was strikingly similar to what has been described for *znfx-1* mutants (Ishidate et al., 2018). Interestingly, loss of ZNFX-1 revealed a remarkable change in 22G RNA distribution over the length of the gene body of target loci: 22G RNAs were mostly lost from their 3’ end, whereas 22G RNAs from the 5’ part of these genes increased. Hence, we probed for 22G RNA coverage on the gene bodies of the various gene sets previously mentioned, using a metagene analysis as employed by Ishidate et al., 2018. This revealed that mutator target genes overall lost 22G RNAs mostly from their 5’ regions in *pid-2* mutants (Figure 3C), a finding opposite to what has been described for *znfx-1* mutants. This was not observed for other gene sets, with the possible exception of CSR-1 targets (Figure 3C). We conclude that PID-2 affects 22G RNA production from many loci, including many previously defined mutator targets, and that these can either gain or lose 22G RNAs. In addition, PID-2 is required to specifically maintain 22G RNA production from the 5’ parts of the target transcripts, suggesting a role in RdRP processivity.

**Figure 3.**
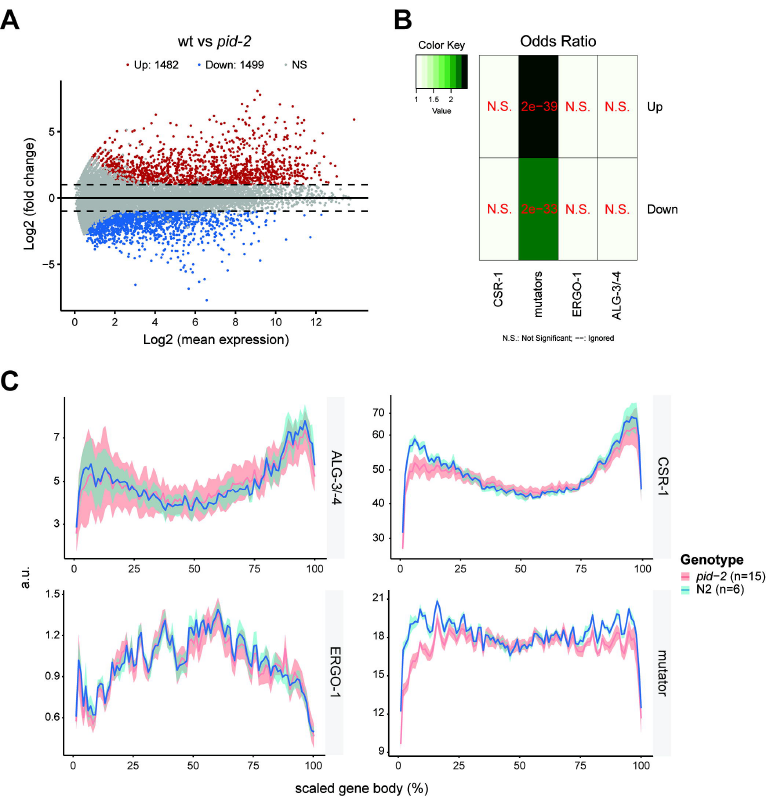
Loss of PID-2 causes a disbalance in 22G RNA populations. **A)** MA-plot of log_2_ fold changes (Y-axis) versus the mean of normalized counts of 22G RNAs (X-axis) for *pid-2* mutants, compared to wild type. Red dots: genes with adjusted p-value < 0.05 and fold-change > 2. Blue dots: genes with adjusted p-value < 0.05 and fold-change < −2. **B)** Heatmap displaying overlap significance between different gene sets and genes that either up- or down-regulated in *pid-2* mutants. Significance was tested with Fisher’s exact test. Colour scheme represents the odds ratio of overlaps representing the strength of association. **C)** Cumulative 22G coverage along the gene body of different sets of genes. Values represent 22G coverage normalized to the total coverage of the gene, for wild type (N2) and *pid-2* mutants. Gene sets are previously defined 22G RNA target sub-types: ALG-3/-4 (Almeida et al., 2019a), CSR-1 (Conine et al., 2013), ERGO-1 (Almeida et al., 2019a), mutator (Phillips et al., 2014). The lines represent the average of biological replicates, whereas the shading represents the standard deviation of biological replicates. a.u.: arbitrary units.

### PID-2 interacts with two eTudor-domain proteins: PID-4 and PID-5

To define the molecular environment of PID-2, we performed immunoprecipitation (IP) followed by quantitative label-free mass spectrometry (MS) on gravid adults, using both the transgenic line expressing the rescuing, C-terminally tagged PID-2::eGFP fusion protein (*xfIs145*), as well as a polyclonal antibody, that we raised against the endogenous protein (Figure 4A, B). In addition, we also performed this experiment with an N-terminally 3xFLAG-tagged PID-2 fusion protein expressed from two independent, randomly inserted transgenes generated using the miniMos procedure (Frøkjær-Jensen et al., 2014) (Figure S4A, B). In all cases, we could reproducibly pull down PID-2, indicating that the IP was working well. In addition to PID-2, we consistently identified two non-characterized proteins: W03G9.2 and Y45G5AM.2, even if enrichment of the latter did not reach our significance cutoff in the PID-2::eGFP IP. We named these proteins PID-4 and PID-5, respectively (Figure 4A, B; S4A, B). Both PID-4 and PID-5 are predicted to have an extended Tudor (eTudor) domain, as found by analysis with HHpred (Zimmermann et al., 2018). These domains are known to bind symmetrically dimethylated arginines, which involves a set of four conserved aromatic residues that form a so-called aromatic cage, and a characteristic acidic amino acid (Gan et al., 2019). Neither the aromatic cage, nor the acidic residue are found in the PID-4 and PID-5 eTudor domains (Figure S4C), suggesting that PID-4 and PID-5 do not bind dimethylated arginines. Interestingly, PRMT-5, the enzyme responsible for symmetric dimethylation of arginine, displayed some enrichment in PID-2::eGFP IPs, but did not reach significance (Figure 4A). In addition to an eTudor domain, PID-5 has an X-prolyl aminopeptidase domain, which is very similar to APP-1 (Figure 4C; S4D). APP-1 is a strongly conserved enzyme, found from yeast to human, that cleaves the most N-terminal amino acid from a polypeptide, provided that the second amino acid is a proline (Iyer et al., 2015; Laurent et al., 2001). The PID-5 X-prolyl aminopeptidase domain is likely catalytically inactive, as the residues required to coordinate the two Zn^2+^ ions are not conserved (Figure S4D). Interestingly, in the PID-2 IP-MS experiments we detected APP-1 as enriched (Figure 4A, B; Figure S4A, B), even though its enrichment never reached our stringent significance cut-off. Since APP-1 itself dimerizes (Iyer et al., 2015), this could reflect the presence of PID-5:APP-1 heterodimers (also see below).

**Figure 4.**
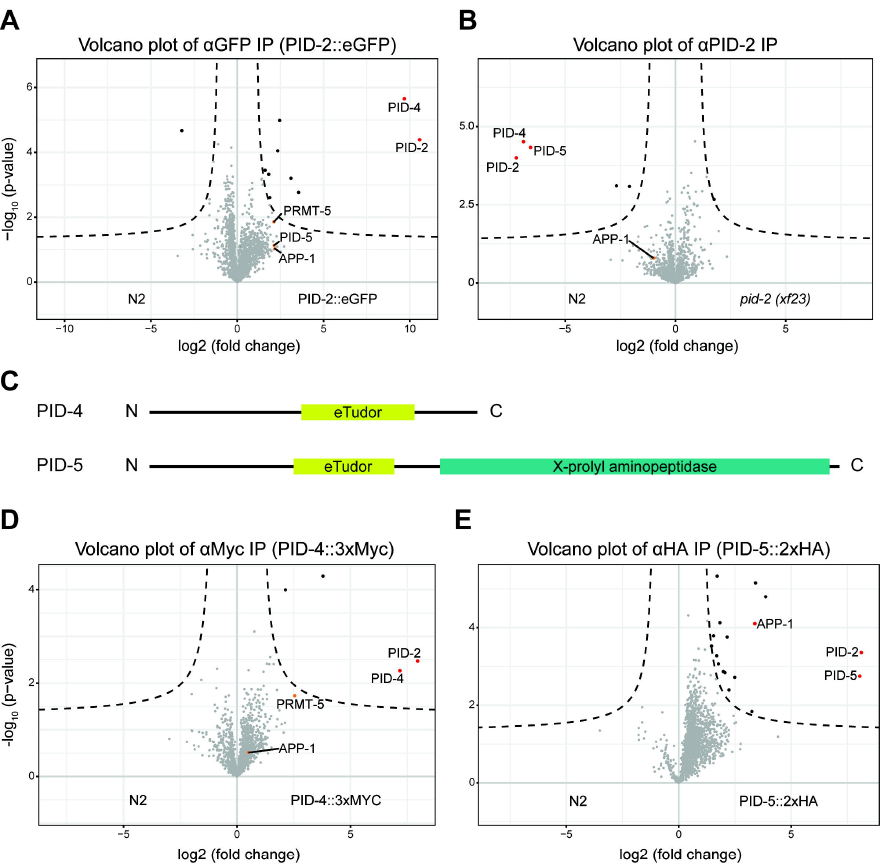
IP-MS on PID-2 identifies two novel interacting proteins, PID-4 and PID-5, and reveals the existence of two distinct complexes. **A)** Volcano plot representing the enrichment of proteins interacting with PID-2, isolated by immunoprecipitation of eGFP::PID-2 protein, followed by quantitative label-free mass spectrometry. As control, eGFP IPs were performed on protein extracts from wild-type, non-transgenic animals. For each IP-MS experiment, quadruplicates were measured and analysed. The dashed line reflects a significance threshold of p-value < 0.05 at two-fold enrichment. **B)** Volcano plot, as described in **A**. In this experiment, the endogenous PID-2 protein was immunoprecipitated, and *pid-2(xf23)* mutant protein extracts were used as control. **C)** Schematic representation of the predicted domain structure of PID-4 and PID-5. The eTudor domains were identified using HHpred (Zimmermann et al., 2018). The X-prolyl aminopeptidase domain was identified using BLAST (https://blast.ncbi.nlm.nih.gov/Blast.cgi). **D)** Volcano plot, as described in **A**, representing the enrichment of proteins interacting with PID-4, by immunoprecipitating endogenously tagged PID-4 protein. IPs from protein extracts from wild-type, non-tagged animals were used as control. **E)** Volcano plot, as described in **A**, representing the enrichment of proteins interacting with PID-5, by immunoprecipitating endogenously tagged PID-5 protein. IPs from protein extracts from wild-type, non-tagged animals were used as control.

We tagged both PID-4 and PID-5 endogenously with an epitope and with a fluorescent tag (Figure S4E, F), in order to perform IP-MS experiments and to investigate their expression. IP-MS on either PID-4 (Figure 4D) or PID-5 (Figure 4E) enriched for PID-2, consistent with their enrichment in PID-2 IPs. In addition, PRMT-5 and APP-1 were enriched in PID-4 and PID-5 IPs respectively (Figure 4D, E), lending support to the detection of APP-1 and PRMT-5 in the above mentioned PID-2 IPs. We did not retrieve PID-5 in PID-4 IPs, or *vice versa*, indicating that PID-4 and PID-5 do not simultaneously interact with PID-2. Consistent with this finding, IP-MS on PID-2 in *pid-4* and *pid-5* mutant backgrounds (Figure S4E, F) still retrieved PID-5 and PID-4, respectively (Figure S4G, H). Collectively, these data identify PID-4 and PID-5 as robust PID-2 interacting proteins. Additionally, APP-1 may interact with the PID-2 complex via PID-5, and PRMT-5 via PID-4.

### PID-4 and PID-5 are partially redundant

Mutant strains lacking PID-4 or PID-5 did not show obvious developmental defects. Given that *pid-2* mutants have defects in the silencing of the 21U sensor, we next investigated the expression of the 21U sensor in *pid-4* and *pid-5* mutants. We introduced a 21U sensor, either active or silenced via RNAe, into *pid-4* and *pid-5* mutant backgrounds. Independent of the initial status of the 21U sensor, both *pid-4* as well as *pid-5* mutants were silencing-proficient (Figure S1E). We hypothesized that PID-4 and PID-5 could be redundant, given that their respective eTudor domains are very similar and could have interchangeable roles (Figure S4C). A *pid-4;pid-5* double mutant strain indeed revealed redundancy, as these double mutants fail to silence a 21U sensor when it is not under RNAe (Figure 5A), with expression levels that are very similar to those found in *pid-2* mutants (Figure S1E). A 21U sensor under control of RNAe was not reactivated (Figure S1E).

**Figure 5.**
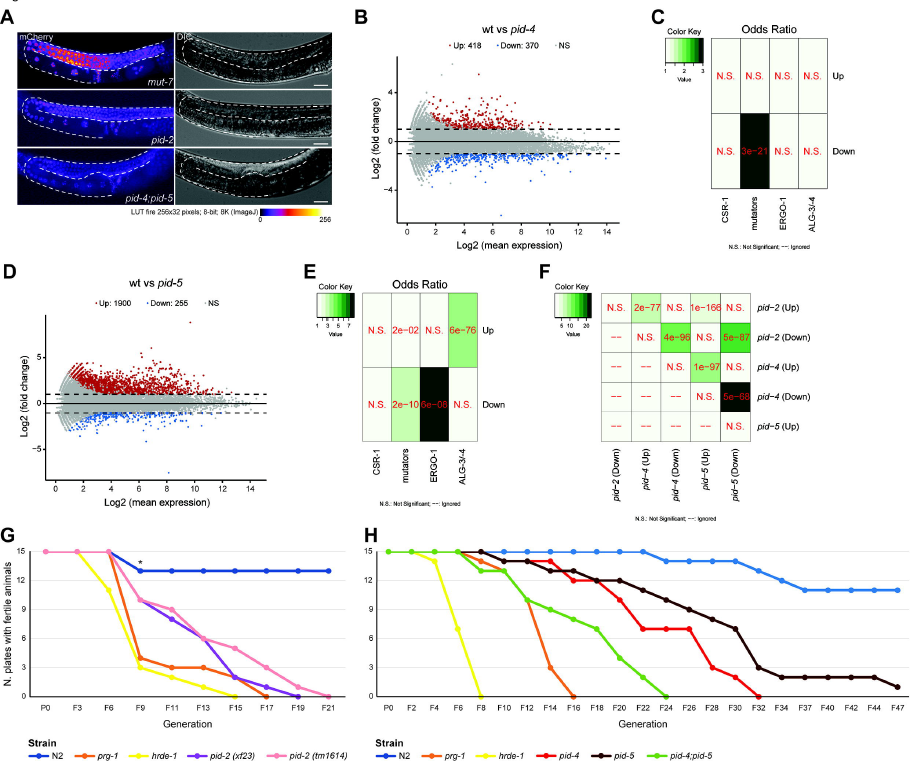
PID-2 and its interactors, PID-4 and PID-5, are required for maintenance of an immortal germline. **A)** Expression of the 21U sensor in the indicated genetic backgrounds. Gonads are outlined by a dashed line. The mCherry signal is represented in pseudo-colours [LUT fire (ImageJ)] to reflect differences in the intensity of the signal. The panels on the right are DIC images of the respective animals. Scale bar: 25 μm. **B)** MA-plot of log_2_ fold changes (on the Y-axis) versus the mean of normalized counts of 22G RNAs (on the X-axis) for *pid-4* mutants, compared to wild type. Red dots: genes with adjusted p-value < 0.05 and fold-change > 2. Blue dots: genes with adjusted p-value < 0.05 and fold-change < −2. **C)** Heatmap displaying overlap significance between different gene sets and genes that either up- or down-regulated in *pid-4* mutants. Significance was tested with Fisher’s exact test. Colour scheme represents the odds ratio of overlaps representing the strength of association. **D)** MA-plot of log_2_ fold changes (on the Y-axis) versus the mean of normalized counts of 22G RNAs (on the X-axis) for *pid-5* mutants, compared to wild type. Red dots: genes with adjusted p-value < 0.05 and fold-change > 2. Blue dots: genes with adjusted p-value < 0.05 and fold-change < −2. **E)** Heatmap displaying overlap significance between different gene sets and genes that either up- or down-regulated in *pid-5* mutants. Significance was tested with Fisher’s exact test. Colour scheme represents the odds ratio of overlaps representing the strength of association. **F)** Heatmap displaying overlap significance between genes that either up- or down-regulated in *pid-2, pid-4* and *pid-5* mutants. Significance was tested with Fisher’s exact test. Colour scheme represents the odds ratio of overlaps representing the strength of association. **G, H)** Line plots representing fertility over generations for the indicated genetic backgrounds at 25°C. Six L2-L3 larvae for each of the indicated backgrounds were hand-picked to a fresh plate every four or six days, counting as two or three generations, respectively, until no larvae were present on the plate to be picked. *: two plates of N2 were contaminated and were excluded from the assay.

We also performed small RNA sequencing on *pid-4* and *pid-5* single mutants, in an attempt to reveal potentially subtle effects of the individual mutations. First, the 21U sensor is normally targeted by 22G RNAs in both mutants (Figure S5A), as it might have been expected from the 21U sensor activity analysis. Moreover, we could not detect striking alterations in the bulk abundance of any of the small RNA classes in *pid-4* and *pid-5* mutants, although 26G RNAs appeared to be slightly, but significantly reduced in *pid-4* mutants (Figure S5B). These data suggest that *pid-4* and *pid-5* mutations on their own have little effect on small RNAs. However, when we looked at 22G RNA profiles in more detail, we did find significant effects of both mutations individually. Loss of PID-4, like loss of PID-2, resulted in genes losing or gaining 22G RNAs (Figure 5B). However, fewer genes were affected (370 genes lost, and 418 genes gained 22G RNAs) in *pid-4* mutants (Figure 5B; Table S2). The genes that lost 22G RNAs in *pid-4* mutants were strongly enriched for mutator targets (Figure 5C), indicating that PID-4 may mainly act to promote 22G RNA production from these genes. The genes that gained 22G RNAs did not show enrichment for any particular gene set we analysed (Figure 5C). Loss of PID-5 showed a strongly asymmetric effect: 255 genes lost, while 1900 genes gained 22G RNAs (Figure 5D; Table S3). The genes that lost 22G RNAs were again enriched for mutator targets, and also for ERGO-1 targets (Figure 5E). Additionally, the genes that gained 22G RNAs in *pid-5* mutants were enriched for ALG-3/-4 target genes (Figure 5E), something that could also be seen on the bulk analysis of subsets of 22G RNAs, and was also detectable at very similar levels in *prg-1* mutants (Figure S5C). When compared to the genes affected by *pid-2*, both *pid-4* and *pid-5* affected genes displayed a striking overlap, both in terms of gene identity, as well as in direction of the detected effect (gain or loss of 22G RNAs; Figure 5F). We did not detect the loss of 22G RNAs specifically from the 5’ or 3’ portions of mutator targets in these single mutant data-sets, but ALG-3/-4 and CSR-1 targets did show differential 5’ or 3’ effects in *pid-4* mutants (Figure S6A, B). These data suggest that PID-4 and PID-5 have partially redundant roles in stimulating 22G RNA production from mutator target genes, while PID-5 has an additional repressive effect on 22G RNA production, which is most noticeable on ALG-3/-4 target genes.

### PID-2, PID-4 and PID-5 are required for an immortal germline

As mentioned earlier, *prg-1* mutants do not show massive transposon reactivation nor acute sterility, however their fertility gradually diminishes over generations. This so-called mortal germline phenotype (Mrt) is not only a characteristic of *prg-1* mutants (Simon et al., 2014), but also of mutants for other factors participating in the RNAe machinery, such as *hrde-1*, *nrde-1/-2/-4* (Buckley et al., 2012), the H3K4 methyltransferase *set-2* (Xiao et al., 2011), the H3K9 methyltransferase homolog *set-32* (Spracklin et al., 2017), and two factors involved in RNAi inheritance, WAGO-4 (Xu et al., 2018) and ZNFX-1 (Wan et al., 2018). Therefore, we tested whether *pid-2*, *pid-4*, *pid-5* or *pid-4;pid-5* mutants showed a Mrt phenotype. As expected, both *prg-1* and *hrde-1* mutants produced very little offspring already after few generations and eventually became sterile, between 6 and 14 generations, whereas the large majority of wild-type worms did not become sterile, even after 47 generations. Also, *pid-2* as well as *pid-4;pid-5* mutants became sterile after 8-9 generations, but the Mrt phenotype was only fully established after 20-24 generations. Strikingly, also *pid-4* and *pid-5* single mutants showed a Mrt phenotype, even though it became apparent only very slowly. However, they did eventually become completely sterile after roughly 30 generations (Figure 5G, H). We note that the presented data tends to under-estimate the effect of the mutations, as we noticed that the numbers of offspring produced by the various mutants already dropped after a few generations, and this is not reflected in the data.

### PID-2, PID-4 and PID-5 localize to perinuclear granules in germ cells

We performed confocal microscopy to investigate the expression pattern and localization of PID-2, tagged with eGFP at its C-terminus (*xfIs145*; Figure S1A). We imaged L4 larvae, as the expression levels of PID-2, as well as of PID-4 and PID-5 (see below), were rather low and most clearly detected in the pachytene stage of the meiotic region, which is most extended at the L4 stage. PID-2::eGFP localized to perinuclear granules. As shown by the colocalization with PGL-1 (Kawasaki et al., 1998), PID-2 foci were adjacent to P granules (Figure 6A). Z granules, marked by ZNFX-1, have been recently described to be juxtaposed to P granules, and involved in transmitting genetic information to the next generation via the oocytes (Ishidate et al., 2018; Wan et al., 2018). As we observed a role for PID-2 in ensuring an efficient silencing by maternally inherited 21U RNAs and in stably maintaining RNAe, PID-2 may well localize to Z granules. The typical distance between PID-2 and PGL-1 foci (Figure 6F), and ZNFX-1 and PGL-1 foci (see below and Figure S7B) closely matched each other, consistent with this idea. Unfortunately, direct assessment of co-localization between ZNFX-1 and PID-2 was thus far hampered by the very close linkage between the PID-2::eGFP transgene and an available endogenously tagged ZNFX-1 allele (Wan et al., 2018). However, Wan et al. (manuscript submitted in parallel to ours) have data that clearly place PID-2 (named ZAP-1 by Wan et al.) in Z granules.

**Figure 6.**
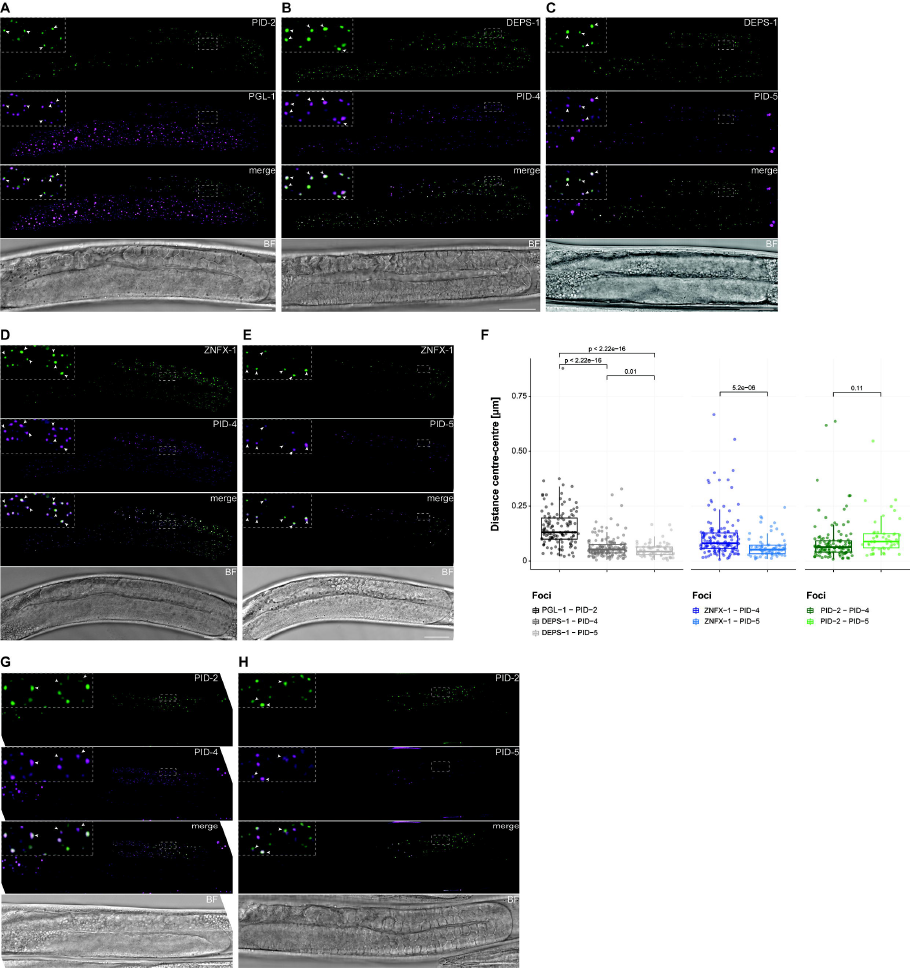
PID-2 is expressed in Z granules, whereas PID-4 and PID-5 localize to P granules, and their expression partially overlaps. **A-C)** Expression of PID-2::eGFP (*xfIs145*; **A**), PID-4::mTagRFP-T (**B**) and PID-5::mTagRFP-T (**C**) in perinuclear granules in the germline. PGL-1::mTagRFP-T (**A**) and DEPS-1::GFP (**B, C**) mark P granules. The indicated dashed boxes reflect zoom-ins on specific nuclei to better visualize the granules, and their overlaps. One L4 gonad is shown for each animal. Note that most of the L4 gonad is in pachytene stage. Scale bar: 25 µm. **D, E)** Expression of PID-4::mTagRFP-T (**D**) and PID-5::mTagRFP-T (**E**) together with 3xFLAG::GFP::ZNFX-1, a Z granule marker. The indicated dashed boxes reflect zoom-ins on specific nuclei to better visualize the granules, and their overlaps. One L4 gonad is shown for each animal. Note that most of the L4 gonad is in pachytene stage. Scale bar: 25 µm. **F)** Box-plots representing the distance (µm) between the centres of two fluorescent signals from the indicated fusion proteins, as represented in panels **A-E, G, H**. The distance between each pair of fluorescent proteins is represented by a dot. The median is represented by a line. The 75^th^ and 25^th^ percentile are represented by the upper and lower lines, respectively. P-values were calculated using a t-test (two-tailed). **G, H)** Expression of PID-2::eGFP (*xfIs145*) together with either PID-4::mTagRFP-T (**G**) or PID-5::mTagRFP-T (**H**). The indicated dashed boxes reflect zoom-ins on specific nuclei to better visualize the granules, and their overlaps. One L4 gonad is shown for each animal. Note that most of the L4 gonad is in pachytene stage. Scale bar = 25 µm.

PID-4 and PID-5 were both endogenously tagged at the C-terminus with mTagRFP-T (Figure S4E, F). We found that PID-4 and PID-5 were also specifically expressed in germ cells and localized to perinuclear granules as well. PID-5 was detectable in foci around relatively few nuclei at the pachytene stage, whereas PID-4 was found throughout the gonad, with stronger expression at the pachytene stage (Figure 6B-E, G, H). PID-4 and PID-5 colocalized to a large extent with the P granule marker DEPS-1 (Spike et al., 2008) (Figure 6B, C). Moreover, the typical distance between PID-4/-5 foci and DEPS-1 was significantly shorter than between PID-2 and PGL-1 (Figure 6F). Interestingly, PID-4 and PID-5 were also relatively close to ZNFX-1 foci (Figure 6D-F). Even if PID-4 displayed a tendency to be further away from Z granules than PID-5, both PID-4 and PID-5 were significantly closer to Z granules than P granules were (Figure 6F). Finally, PID-4/PID-5 distances to Z granules were very similar to the separation between PID-4/PID-5 and PID-2 (Figure 6F-H). We conclude that PID-2 on the one hand, and PID-4 and PID-5 on the other hand, display distinct subcellular localization in discrete perinuclear foci, but also that these foci do display significant overlaps. While PID-2 is in Z granules, assignment of PID-4 and PID-5 to distinct, known perinuclear foci is more difficult, as they form foci that are very close to, yet distinct from both P and Z granules.

### PID-2, PID-4 and PID-5 affect Z granules formation

Finally, we checked whether loss of PID-2, PID-4 or PID-5 affected P or Z granule appearance. Localization of PGL-1 in *pid-2* mutants strongly resembled that of wild-type animals (Figure 7A, B). In contrast, loss of PID-2 did affect Z granules (Figure 7B). First, we noticed the appearance of relatively large Z granules in *pid-2* mutants. To quantify this, we measured the surface of Z granules, and compared these to the area of Z granules in a wild-type strain. Even though many Z granules were similarly sized in both genetic backgrounds, Z granules indeed displayed a tendency to be larger in *pid-2* mutants than in wild-type animals (Figure S7A). In addition, we counted the number of P and Z granules, to determine their ratio. This revealed a significant loss of Z granules compared to P granules in *pid-2* mutants (Figure 7F). Distance measurements showed that Z granules did remain distinct from P granules (Figure S7B). The distance between Z and P granules tended to be slightly longer in *pid-2* mutants (Figure S7B), but this could be a result of the tendency of Z granules to be larger (Figure S7A). We also checked PID-4 and PID-5 localization in *pid-2* mutants. This revealed that PID-4 foci were affected in *pid-2* mutants. PID-4 foci can still be observed, but they were fewer than in wild-type animals and the intensity of the signal was lower, indicating that PID-4 expression was reduced in absence of PID-2, and/or that PID-4 localization was affected (Figure 7G, S7C). PID-5 foci were not affected by loss of PID-2 (Figure 7H, S7D). Interestingly, the remaining PID-4 foci in *pid-2* mutants were further away from ZNFX-1 compared to wild type (Figure 7I). This could be a result of increased Z granule size, even though for PID-5 this effect was not detected (Figure 7I).

**Figure 7.**
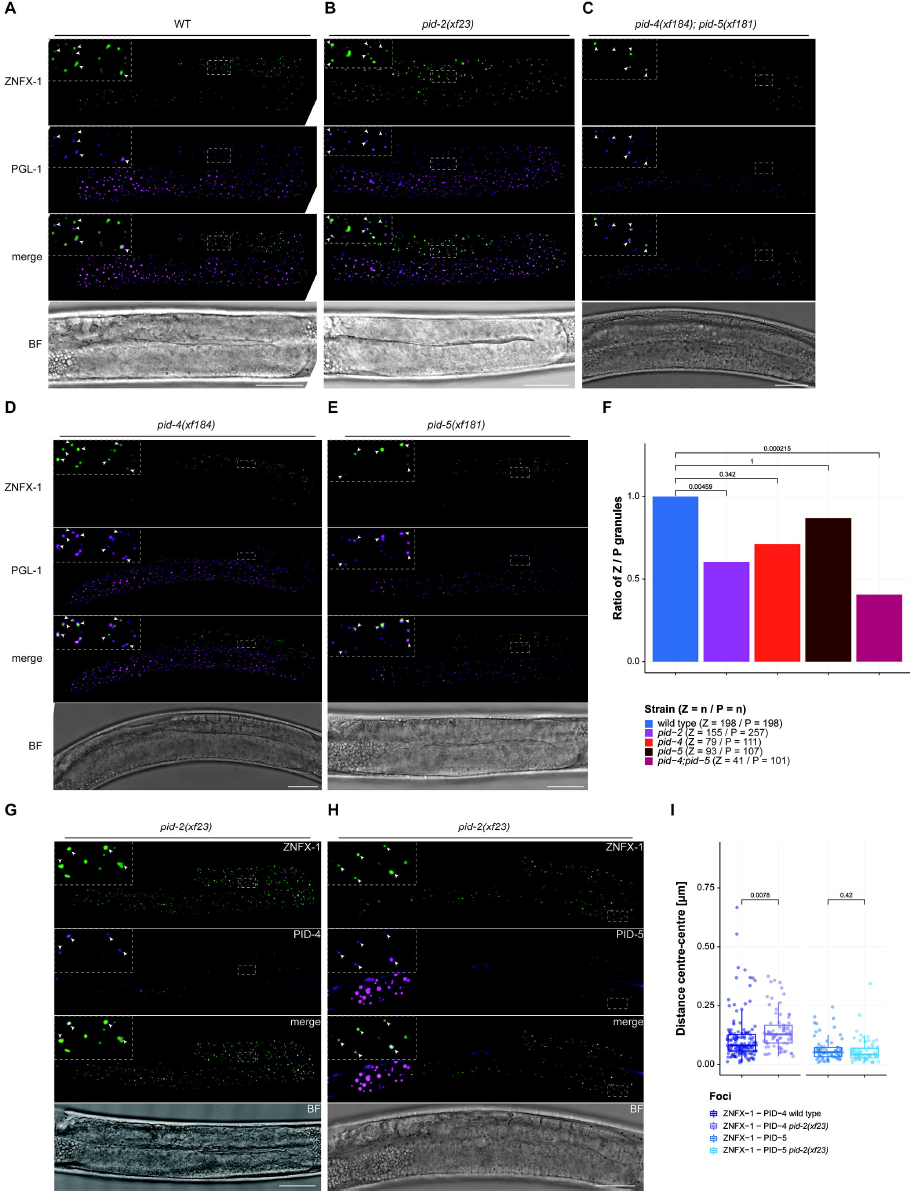
Z granules are affected by PID-2, PID-4 and PID-5. **A-E)** Expression of 3xFLAG::GFP::ZNFX-1 and PGL-1::mTagRFP-T in a wild type (**A**), *pid-2* (**B**), *pid-4;pid-5* (**C**), *pid-4* (**D**), and *pid-5* (**E**) mutant backgrounds. The indicated dashed boxes reflect zoom-ins on specific nuclei to better visualize the granules, and their overlaps. One L4 gonad is shown. Scale bar: 25 µm. **F)** Quantification of the ratio Z / P granules in wild type, *pid-2*, *pid-4/-5*, *pid-4* and *pid-5* mutant backgrounds. The number of granules is indicated in brackets, next to the genotype. P-values were calculated using a two-proportion z-test and adjusted for multiple comparisons with the Holm method. **G, H)** Expression of PID-4::mTagRFP-T (**G**) and PID-5::mTagRFP-T (**H**) together with 3xFLAG::GFP::ZNFX-1, a Z granule marker, in a *pid-2* mutant background. The indicated dashed boxes reflect zoom-ins on specific nuclei to better visualize the granules, and their overlaps. One L4 gonad is shown for each animal. Scale bar: 25 µm. **I)** Box-plots representing the distance (µm) between the centres of two fluorescent signals from the indicated fusion proteins, as represented in panels **G, H** and in Figure 6D, E. Please note that the wild-type distance measurements are the same represented in Figure 6F. The distance between each pair of fluorescent proteins is represented by a dot. The median is represented by a line. The 75^th^ and 25^th^ percentile are represented by the upper and lower lines, respectively. P-values were calculated using a t-test (two-tailed).

Finally, we checked whether loss of PID-4 and/or PID-5 affected Z or P granules. Like in *pid-2* mutants, P granules were not detectably affected, in either single or double mutants (Figure 7C, D, E). However, ZNFX-1 was affected in *pid-4;pid-5* double mutants (Figure 7C). In particular, the number of Z granules were found to be lower (Figure 7C, F), an effect that also tended to appear in *pid-4* and *pid-5* single mutants, but did not reach significance (Figure 7D, E, F). An effect on Z granule size could not be detected (Figure S7A). To check whether the observed effects on Z granules could stem from effects on ZNFX-1 stability, we performed Western blot analysis on endogenously tagged ZNFX-1 in the different mutant backgrounds (Figure S7E). This did not reveal major changes in ZNFX-1 levels. Everything considered, we conclude that PID-2, PID-4 and PID-5 affect steady-state Z granule appearance, while not affecting P granules visibly. Additionally, loss of PID-2 versus loss of PID-4/-5 do not perfectly phenocopy each other at this level, indicating that some PID-2 function remains in *pid-4;pid-5* double mutants, and/or that some PID-4/-5 function remains in *pid-2* mutants.

## Discussion

We describe the identification and characterization of three novel proteins, PID-2, PID-4 and PID-5, that are required to establish silencing on a 21U RNA target, and for normal 22G RNA homeostasis. They localize to distinct germ cell specific perinuclear granules, and affect the morphology of Z granules. We will discuss below potential modes of action of these proteins, and in addition touch upon some evolutionary aspects.

### PID-2 has an essential role in *de novo* silencing mediated by maternally inherited 21U RNAs

We have shown that maternally provided 21U RNAs are required and sufficient for *de novo* target silencing by RNAe, and that they absolutely require PID-2 to do so. In contrast, PID-2 only mildly affects silencing that is already established. Two important conclusions can be drawn from this. First, initiation and maintenance of 21U RNA-induced silencing have different requirements. One possibility we envisage is that PID-2 may act to ‘boost’ the 22G RNA response, and that this ‘boost’ is essential to activate RNAe, while it merely helps to stabilize, and is not essential for established RNAe. How PID-2 may do this will be discussed in the following paragraph. Second, these results identify embryogenesis as an important developmental period for establishment of 21U RNA-mediated silencing. It seems likely that PRG-1 most effectively establishes RNAe in the primordial germ cells. Whether perfect base-pairing between a 21U RNA and its target is required to establish RNAe remains unclear for now, but the fact that 21U RNAs are destabilized in *henn-1* mutants specifically during embryogenesis (Billi et al., 2012; Montgomery et al., 2012) implies that perfect 21U RNA-target interactions are at least prevalent at this stage. Seen from this perspective, PID-2 plays an important role in interpreting the genetic information inherited from the parents and in initiating long-lasting silencing effects in the embryo.

The idea that maternal Piwi activity is particularly important in the PGCs is not unique to *C. elegans*. Also in *Drosophila* maternal impact of Piwi proteins and piRNAs have been described (Akkouche et al., 2017; Siddiqui et al., 2012), and in zebrafish we found significant maternal effects on piRNA populations that differ among strains (Kaaij et al., 2013). Additionally, in plants strong maternal effects of small RNAs have been described, and in unicellular eukaryotes, like *Tetrahymena* and *Paramecium*, similar mechanisms operate in which parental nuclei produce small RNAs that act in the nuclei generated during mating (Castel & Martienssen, 2013; Malone & Hannon, 2009; van Wolfswinkel & Ketting, 2010). Hence, the concept that small RNAs from the parents prime effects in the offspring appears to be an ancient aspect of these pathways. It may therefore not be surprising that the PGCs in animal embryos may contain specialized small RNA-related mechanisms compared to the adult germline. Establishment of RNAe, in which PID-2 plays an important role, may be one such unique property of the PGCs in *C. elegans*.

### What is the molecular mechanism behind the observed phenotypes?

We identified PID-2 because it has a role in the 21U RNA pathway, as shown by the defects in silencing of the 21U RNA sensor. Considering our finding that PID-2 affects 22G RNAs targeting the 21U sensor, we consider a potential function of PID-2 between PRG-1 target recognition and RdRP activity. PID-2 is not essential for silencing, since significant silencing of a 21U sensor can be achieved in *pid-2* mutants. Hence, it seems more likely that PID-2 regulates factors that in turn execute the silencing response. We also note that PID-2 is not specific to the PRG-1 pathway, as *pid-2* mutants do show effects, albeit small, on overall 22G RNAs, including those of the CSR-1 pathway, as well as relatively strong effects on 26G RNAs. Possibly, PID-2 could have a general function in the regulation of RdRP activity, and thus affect both 22G and 26G RNAs. The fact that in *pid-2* mutants, mutator target genes lose 22G RNAs preferentially within their 5’ regions also supports this idea: PID-2 could be involved in the processivity of the RdRP enzyme RRF-1. We note that this effect is opposite to what has been observed for *znfx-1* mutants, that lose 22G RNAs preferentially within the 3’ end of target genes (Ishidate et al., 2018). In biological processes, stable states are typically achieved by applying opposite forces; ZNFX-1 and PID-2 may represent two such oppositely acting mechanisms to ensure stable RdRP activity.

However, there are also other possibilities. With the available data, it is very difficult to dissect primary from secondary effects. Hence, the effect of PID-2 on both 22G and 26G RNAs could be indirect. Indeed, loss of 26G RNAs has also been observed in *mut-16* mutants (Zhang et al., 2011), suggesting that 26G RNA biogenesis may be coupled to 22G RNA biogenesis. Such interactions could also exist between RNAe initiation and maintenance mechanisms. As we noticed in our analysis of Tc1 activity and RNAe maintenance in *pid-2* versus *pid-2;prg-1* double mutants, compensatory mechanisms may exist, meaning that certain endogenous small RNA pathways may become stronger when another gets weaker. Along these lines, we note that the 22G RNA levels of the 21U sensor are higher in *prg-1* mutants compared to wild type (Figure 2), further strengthening this idea. Since *pid-2*, *pid-4* and *pid-5* mutants display both gain and loss of 22G RNAs from specific loci, a role for these factors in balancing the activities of different 22G RNA pathways may seem more likely than one specifically in promoting RdRP activity. Finally, on top of these mechanistic complications, developmental defects may further convolute phenotypes. We noticed feminization and masculinization phenotypes in our experiments, and a specific 21U RNAs has been shown to act in sex determination (Tang et al., 2018). Such effects could play a role in, for instance, the upregulation of 22G RNAs from ALG-3/-4 targets in *pid-5* mutants, as these targets are enriched for spermatogenesis-specific functions. Clearly, biochemical experiments will be required to arrive at a defined molecular functions of the newly identified, but also already known small RNA pathway components.

### A function for the eTudor domains of PID-4 and PID-5

PID-4 and PID-5 were identified as robust PID-2 interactors. Both proteins contain an eTudor domain, and mutants lacking both PID-4 and PID-5 behave very similar to *pid-2* mutants, suggesting that PID-2 acts through PID-4 and PID-5. We do not know how the PID-4/-5 interactions with PID-2 are mediated. Given that PID-4 and PID-5 cannot simultaneously interact with PID-2, and that the PID-4 and PID-5 eTudor domains are similar, it is a good possibility that these eTudor domains bind to PID-2. Many eTudor domains recognize and bind symmetrically di-methylated arginine or lysine residues of partner proteins (Gan et al., 2019; Pek et al., 2012). However, sequence alignments suggest that the eTudor domains of PID-4 and PID-5 may not bind di-methylated arginines. Curiously, our IP-MS experiments on PID-2 and PID-4 revealed mild enrichment for PRMT-5, an enzyme known to symmetrically dimethylate arginines (Pek et al., 2012; Siomi et al., 2010). It is thus possible that PID-2, via PID-4, brings PRMT-5 activity into the small RNA systems of *C. elegans*, in order to modify arginines on other small RNA pathway components. Such potential substrates for PRMT-5 could be PRG-1, as well as HRDE-1 or one of the RdRPs, which all contain several RG motifs. Among the RdRPs, especially RRF-3 has a relatively large number of such motifs (10).

### Potential roles for PID-2, PID-4 and PID-5 in granule dynamics

We found that PID-2 on the one hand, and PID-4 and PID-5 on the other hand, are present in distinct perinuclear granules. PID-4 and PID-5 foci overlap with P granules, but also show significant overlap with Z granules. PID-2 instead forms foci that are clearly distinct from P granules, but adjacent to them, and Wan et al. (accompanying manuscript) could show that PID-2 (named ZAP-1 in their study) is indeed present in Z granules. Whether PID-4 and PID-5 are in P or Z granules can currently not be resolved, as we find close proximity to both. Possibly, they may be present in another, yet unknown, kind of perinuclear compartment which would be closely associated to both P and Z granules. Interestingly, PID-2 affects PID-4, and to a lesser extent also PID-5 localization, even though they appear to be in different granules. The simple fact that several proteins acting at different steps in the mutator pathway are found enriched in different phase-separated structures (Batista et al., 2008; Claycomb et al., 2009; Ishidate et al., 2018; Phillips et al., 2012; Updike & Strome, 2010; Wan et al., 2018; Wang & Reinke, 2008) indicates that exchange of molecules between different granules will be required to ensure an efficient silencing pathway. Yet, not much is known about such trafficking between adjacent phase-separated structures, not only as a general mechanism, but also not in relation to small RNA pathways. Our study identifies a set of three novel proteins that are excellent candidates to act in such a process, providing important first handles to study this. We note that our localization studies are based on proteins tagged with fluorophores, and we can currently not exclude that these tags interfere with localization. However, the fact that all three identified PID proteins affect Z granule formation, without affecting ZNFX-1 protein levels, strongly supports a role in Z granule homeostasis.

### A role for X-prolyl aminopeptidase activity in 22G and 26G RNA pathways?

Next to an eTudor domain, PID-5 has an X-prolyl aminopeptidase domain. Based on the loss of key catalytic residues, it is likely catalytically inactive. What could the function of such a protease-like domain be? Potentially, PID-5 could use this domain to bind and lock proteins that carry a proline at position 2, without cleaving the most N-terminal amino acid. This would require a stable association of this catalytically dead X-prolyl aminopeptidase domain. We are not aware of studies assessing the stability of substrate-enzyme interactions of catalytically dead X-prolyl aminopeptidases. Another exciting hypothesis is that PID-5 could use its aminopeptidase domain to dimerize with APP-1. Indeed, the catalytic domain of APP-1 is known to dimerize (Iyer et al., 2015), and we found APP-1 significantly enriched specifically in those IP-MS experiments in which PID-5 was enriched. We envisage that such heterodimerization could have two functions. First, it could bring the enzymatic activity of APP-1 into PID-5 positive granules. Alternatively, PID-5 could inhibit the catalytic activity of APP-1, by outcompeting APP-1 homodimers. In the first model, APP-1 would not be expected to be present in P or Z granules in *pid-5* mutants, whereas in the second model, the localization of APP-1 would be independent of PID-5. We are unfortunately not aware of studies describing whether APP-1 needs to dimerize to be active or not. These questions have to be addressed in future experiments. Another issue that needs to be addressed is the identification of APP-1/PID-5 substrates. WAGO-3, WAGO-4 and WAGO-10 are interesting candidates, as based on the current gene models, they are WAGO proteins with suitable N-termini.

### Evolutionary considerations

The fact that PID-4 and PID-5 bind to PID-2 in a mutually exclusive manner could point at a regulatory interaction between these two proteins. Since PID-5 carries an additional domain, an appealing hypothesis would be that PID-4 could act to modulate the amount of PID-5 that can bind to PID-2. However, both proteins could also have completely independent functions. In this light, the following observation is of interest: The *pid-4* gene is positioned directly next to *app-1* in the *C. elegans* genome. A scenario in which *pid-5* was formed by a gene duplication event, in which *pid-4* and *app-1* became joined together, seems an interesting possibility. Indeed, *pid-5* orthologs are only present in some of the *Caenorhabditis* species (*C. remanei*, *C. brenneri*, *C. briggsae*) (Figure S4D), but not, for instance, in *C. japonica*, which is evolutionary more distant from the above-mentioned species (Kanzaki et al., 2018). However, a *pid-4* ortholog is present in *C. japonica*, and is also located next to *app-1*. This is consistent with the idea that a genomic rearrangement between *pid-4* and *app-1* may have happened in the last common ancestor of *C. elegans*, *C. remanei*, *C. brenneri* and *C. briggsae*, leading to the formation of *pid-5*. This implies that PID-4 may still have a function of its own in *C. elegans*, and likely also in *C. japonica*, consistent with our finding that *pid-4* mutants do have phenotypes on their own.

## Materials and Methods

### Strains maintenance

Worm strains have been grown according to standard laboratory conditions on NGM plates seeded with Escherichia coli OP50 and grown at 20 °C, unless otherwise stated (Brenner, 1974). We used the N2 Bristol strain as wild-type strain. Strains used in this study are listed in the supplemental data.

### Microscopy

20-25 worms have been picked to a drop of M9 (80 µl) on a slide, washed and then fixed with acetone (2 x 80 µl). After acetone has evaporated, worms have been washed 2 x 10 minutes with 80 µl of PBSTriton X100 0,1%. After removing the excess of PBS-Triton X100 0,1%, the worms have been mounted on a coverslip with Fluoroshield™ with DAPI (5 µl) (Art. No. F6057, Sigma).

Alternatively, for live imaging, 20-25 worms have been picked to a drop of M9 (80 µl) on a slide, washed and then 2 µl of 1 M NaN_3_ have been added to paralyze the worms. After removing the excess of M9, a slide prepared with 2% agarose (in water) has been placed on top of the coverslip and worms have been imaged directly.

Images have been acquired either at a Leica DM6000B microscope (objective HC PL FLUOTAR 20x 0.5 dry, Art. No. 11506503, Leica) or at a Leica TCS SP5 STED CW confocal microscope (objective HCX PL APO ’CS 63x / 1.2 water UV, Art. No. 11506280, Leica). Images have then been processed with Leica LAS software and ImageJ. Images representing the expression of PID-2, PID-4, PID-5, and P and Z granules markers have been processed with the Huygens Remote Manager v3.6 and deconvoluted (Huygens Deconvolution, SVI).

For scoring the 21U sensor as active or silenced, we have used a Leica M165FC widefield microscope. The 21U sensor has been scored as: active, if the fluorescence was easily visible with a lower magnification (Plan APO 1.0x, Art. No. 10450028; Leica); faint, if the fluorescence was only visible with a higher magnification (Plan APO 5.0x/0.50 LWD, Art. No. 10447243; Leica); silenced, if no fluorescence was visible. The worms have been later used also for live imaging with a Leica DM6000B microscope as described above.

#### Colocalization analysis

In order to perform colocalization analysis of the fluorescently tagged proteins PID-2, PID-4, PID-5 and of the P and Z granules markers, we used the DiAna plugin of (Fiji Is Just) ImageJ (Gilles et al., 2017; Schindelin et al., 2012). We used the deconvoluted images (Huygens Deconvolution, SVI), which consist of a single z-stack, and analysed the two fluorescence channels of interest at the time. First, we cropped an area containing 1-8 nuclei within the pachytene zone of the gonad arm, adding the area to the ROI manager, to ensure cropping of the same area in both channels being analysed. We then used DiAna_Segment to apply a mask to the images, considering all objects with size from 1 to 2000 pixels and then adjusted the threshold to ensure a faithful segmentation of the images. After segmentation, we performed the analysis using DiAna_Analyse and measured the surface area (µm^2^), the distance between the two signals (µm) and the number of objects present in the cropped area. We then represented the distance between centres as a measure of colocalization. For each couple of fluorescent proteins, we analysed 4-10 images of gonads from individual animals.

### Small RNA sequencing

#### RNA extraction

Synchronized gravid adults have been collected with M9 and fast frozen on dry ice in 250 µl of Worm Lysis Buffer (200 mM NaCl; 100 mM Tris HCl pH=8.5; 50 mM EDTA pH=8; 0,5% SDS). 30 µl of Proteinase K (20 mg/ml; Art. No. 7528.1, Carl Roth) have been added to dissolve the worms for 90 minutes at 65 °C with gentle shaking. Lysate has been centrifuged at maximum speed for 5 minutes at room temperature (RT) and the supernatant was transferred on a Phase Lock Gel tube (Art.No. 2302830, QuantaBio). 750 µl TRIzol LS (Art. No. 10296028, Invitrogen™) have been added per 250 µl of sample and, after homogenization, the samples have been incubated for 5 minutes at RT to allow complete dissociation of the nucleoprotein complex. Then 300 µl of chloroform (Art. No. 288306, Sigma-Aldrich) were added per 750 µl of TRIzol LS and the samples were incubated for 15 minutes at room temperature after mixing. Samples have been centrifuged at 12000 x g for 5 minutes at RT and another round of chloroform extraction has been performed. The aqueous phase has been then transferred to an Eppendorf tube and 500 µl of cold isopropanol was added to precipitate the RNA; samples have been mixed vigorously, incubated at RT for 10 minutes and spun down at maximum speed for at least 10 minutes at 4 °C. The pellet was then washed twice with 1 ml of 75% ethanol and centrifuged for 5 minutes at 7500 x g at 4 °C. The pellet has been dried and diluted in 50 µl of nuclease-free water with gentle shaking for 10 minutes at 42 °C. In order to remove any contamination of genomic DNA, 5 µl of 10X TURBO™ DNase Buffer and 1 µl of TURBO™ DNase (Art. No. AM2238, Invitrogen™) were added to the RNA and incubated at 37 °C for 30 minutes with gentle shaking. The reaction has been stopped by adding 5 µl of 10X TURBO™ DNase Inactivation Reagent. Samples have been centrifuged at 10000 x g for 90 seconds and RNA transferred to a fresh tube. RNA quality has been assessed at Nanodrop and on agarose gel and then samples have been further processed for enrichment of small RNA populations.

#### Library preparation and sequencing

For each strain, three biological replicates have been used for RNA extraction and library preparation. RNA was treated with RppH (RNA 5’ Pyrophosphohydrolase, Art. No. M0356S, New England Biolabs) to dephosphorylate small RNAs and specifically enrich for 22G RNAs, as previously described (Almeida et al., 2019b). For each sample, 1 µg of RNA was incubated for 1 hour at 37 °C with 5 units of RppH and 10X NEB Buffer 2. After dephosphorylation, 500 mM EDTA was added and samples were incubated for 5 minutes at 65°C to stop the RppH treatment and RNA was purified with sodium chloride/isopropanol precipitation. After purification, the recovered total RNA was used for library preparation.

Amplified libraries were purified by running an 8% TBE gel and size-selected for the 146-168bp fraction. Libraries were profiled in a High Sensitivity DNA on a 2100 Bioanalyzer (Agilent technologies) and quantified using the Qubit dsDNA HS Assay Kit, in a Qubit 2.0 Fluorometer (Life technologies).

#### Read procession and mapping

Before mapping to the reference sequences, reads were processed in the following manner: (i) trimming of sequencing adapters with cutadapt v1.9 (-a TGGAATTCTCGGGTGCCAAGG -a AGATCGGAAGAGCACACGTCT -O 5 -m 26 -M 38) (Martin, 2011); (ii) removal of reads with low-quality calls with the FASTX-Toolkit v0.0.14 (fastq_quality_filter -q 20 -p 100 -Q 33); (iii) collapsing of PCR duplicates (custom bash script), making use of the unique molecule identifiers (UMIs) added during library preparation; (iv) trimming of UMIs with seqtk v1.2 (seqtk trimfq -b 4 -e 4); and (v) removal of very short sequences with seqtk v1.2 (seqtk seq -L 15). Read quality was assessed before and after these processing steps with FastQC v0.11.5 (http://www.bioinformatics.babraham.ac.uk/projects/fastqc).

Reads that passed the above filtering steps were mapped to a custom *C. elegans* genome (WBcel235) to which the 21U sensor sequence (Bagijn et al., 2012) was added as an extra contig. The mapping was done with bowtie v0.12.8 (-q –sam –phred33-quals –tryhard –best –strata –chunkmbs 256 -v 0 -M 1) (Langmead et al., 2009). To generate genome browser tracks we used a combination of Bedtools v2.25.0 (genomeCoverageBed -bg -split -scale -ibam -g) (Quinlan & Hall, 2010), to summarize genome coverage normalized to mapped non-structural reads (rRNA/tRNA/snoRNA/snRNA) * 1 million (RPM, Reads Per Million), and bedGraphToBigWig to finally create the bigwig tracks. More detailed information is available in the supplemental data file.

### Transposon excision analysis

For each analysed genotype, mutant worms carrying the *unc-22::Tc1(st136)* insertion were singled into a 6 cm^2^ NGM plate seeded with 100 µl of OP50 and grown at 20 °C. For each genotype, 50 worms have been singled out. Plates were scored for wild type moving worms at three different time points: when the total number of worms per plate was ~50, when the total number of worms per plate was ~100, and when the plate was starved, to which we estimated the total number of worms per plate to be ~1000. Transposition frequencies at each time point were calculated using the following formula: f = -ln [(T - R) / T] / N, where T = total number of plates scored, R = number of plates with revertants, and N = number of worms on the plate. Each time point was considered as a biological replicate. The graph represents two independent experiments and the transposition frequency calculated when the total number of worms per plate was estimated to be ~1000.

### Mortal germline assay

Before starting the experiment, mutants have been outcrossed four times. For the assay, N2 have been used as wild-type strain and the desired mutant strains have been tested. For each strain, 6 L3 worms have been picked onto 15 NGM plates (10 cm^2^) seeded with 300 µl OP50, grown at 25 °C and followed over time. Worms have been picked every 4-6 days, before starvation, and we assumed 2-3 generations, respectively, have passed. Worms have been passed to fresh plates every 4-6 days until all the mutants died.

### Production of PID-2 protein for antibody generation

Information available in the supplemental data file.

### Transgenic lines generation

Information available in the supplemental data file.

### Generation of mutant and endogenously tagged lines using CRISPR/Cas9 technology

#### Cloning

All the sgRNAs have been cloned in the vector p46169 (a gift from John Calarco, (Friedland et al., 2013)), except for Y45G5AM.2_sgRNA7, which has been cloned in the vector pRK2412 (pDD162 backbone, Cas9 deleted with improved sgRNA(F+E) sequence, as described in (Chen et al., 2013)).

#### Generation of mutant lines

Wild type worms have been injected with an injection mix containing 50 ng/µl pJW1259 (encoding for *Peft-3::cas9::tbb-2 3’UTR*, a gift from Jordan Ward (Ward, 2015)), co-injection markers (10 ng/µl pGH8; 5 ng/µl pCFJ104; 2,5 ng/µl pCFJ90) and 30 ng/µl of each of the plasmids encoding for the sgRNAs. We isolated two deletion alleles of *Y45G5AM.2/pid-5* (*xf181* and *xf182*) and two deletion alleles of *W03G9.2/pid-4* (*xf184* and *xf185*). Each allele has been sequenced to pinpoint the exact deletion at nucleotide resolution. The mutant strains have been outcrossed two times against wild type N2 strain to remove any potential off-targets effect of Cas9 and used for further experiments.

#### Generation of endogenously tagged lines

In order to introduce an epitope tag at endogenous loci we used the co-conversion approach as previously described (Arribere et al., 2014). After injections, worms have been kept at 20 °C and F1 offspring with a roller phenotype (*rol-6*) were singled out. The tagged strains have been outcrossed two times against wild type N2 strain to remove any potential off-targets effect of Cas9 and used for further experiments.

To introduce a fluorescent protein at the endogenous locus, we have first used a *unc-58* co-conversion approach (Arribere et al., 2014) to introduce a sequence of 20 nucleotides of *dpy-10* gene that serves as efficient protospacer sequence for subsequent edits, as previously described (Mouridi et al., 2017). The tagged strains were sequenced, and were outcrossed two times against wild type N2 strain to remove any potential off-targets effect of Cas9 and used for further experiments.

Sequences and more detailed information on procedures are available in the supplemental data.

### Protein extraction and immunoprecipitation

Information available in the supplemental data file.

### Mass Spectrometry

Label-free quantitative mass spectrometry has been performed as described in (Almeida et al., 2018). After boiling (see above), the samples were separated on a 4–12% gradient Bis-Tris gel (NuPAGE Bis-Tris gels, 1.0 mm, 10 well; Art. No. NP0321; Life Technologies) in 1X MOPS (NuPAGE 20X MOPS SDS running buffer; Art. No. NP0001; Life Technologies) at 180 V for 10 min, afterwards processed by ingel digest (Bluhm et al., 2019; Shevchenko et al., 2007) and desalted using a C18 StageTip (Rappsilber et al., 2007). The digested peptides were separated on a 25-cm reverse-phase capillary (75 μM inner diameter) packed with Reprosil C18 material (Dr. Maisch) with a 2 h gradient from 2 to 40% Buffer B (see Stage tip purification) with the EASY-nLC 1,000 system (Thermo Scientific). Measurement was done on a Q Exactive Plus mass spectrometer (Thermo Scientific) operated with a Top10 data-dependent MS/MS acquisition method per full scan (Bluhm et al., 2016). The measurements were processed with the MaxQuant software, version 1.5.2.8 (Cox & Mann, 2008) against the Wormbase *C. elegans* database (version of WS265) for quantitation and the Ensemble *E. coli* REL606 database (Version Oct 2018) to filter potential contaminations.

### Protein extraction and Western blot

Information available in the supplemental data file.

### Data availability

Sequencing data is available at NCBI-SRA, BioProject ID PRJNA612883. Mass spectrometry data has been deposited at PRIDE, accession number PXD018402.

## Author Contribution

MP, AMJD and RFK planned experiments. MP generated strains, performed genetics, microscopy and molecular biology. AMJD analysed sequencing data. JS generated strains and performed microscopy. SH generated strains. BFA identified *pid-2* and performed initial analyses. SD and FB performed quantitative mass spectrometry. All authors were involved in discussion of the data. MP and RFK wrote the manuscript with input from all authors.

## Acknowledgements

We thank members of the Ketting lab for stimulating discussions. We thank Walter Bronkhorst and Roberto Orrù for valuable technical assistance. We also thank the IMB Core facilities Media Lab, Microscopy, Bioinformatics, Protein production and Genomics for excellent experimental and technical support. MP was supported by a Boehringer Ingelheim Fonds PhD Fellowship. This work was further supported by grants, KE 1888/1-1 and KE1888/1-2, from the Deutsche Forschungsgemeinschaft (RFK), and by a grant from Fundação para a Ciência e Tecnologia ([FCT]SFRH/BD/51001/2010; B.F.M.A.).

