## Supplemental material for "The IDR-containing protein PID-2 affects Z granules and is required for piRNA-induced silencing in the embryo"

### MATERIALS AND METHODS

#### List of strains used in this study

| Strain | Genotype |
|---|---|
| | wild type N2 |
| RFK507 | *pid-2(xf23) I* |
| RFK345 | *pid-2(tm1614) I* |
| RFK231 | *mjSi22[Pmex-5::mCherry::his-58::21UR-1_as::tbb-2(3'UTR)] I; mut-7(pk204) III; otIs45[Punc119::GFP] V* |
| RFK316 | *mjSi22[Pmex-5::mCherry::his-58::21UR-1_as::tbb-2(3'UTR)] (RNAe) I; prg-1(n4357) I* |
| RFK851 | *mjSi22[Pmex-5::mCherry::his-58::21UR-1_as::tbb-2(3'UTR)] I; prg-1(n4357) I* |
| RFK677 | *pid-2(xf23) I; mjSi22[Pmex-5::mCherry::his-58::21UR-1_as::tbb-2(3'UTR)] I* |
| RFK585 | *pid-2(xf23) I; mjSi22[Pmex-5::mCherry::his-58::21UR-1_as::tbb-2(3'UTR)] (RNAe) I* |
| RFK528 | *pid-2(tm1614) I; mjSi22[Pmex-5::mCherry::his-58::21UR-1_as::tbb-2(3'UTR)] I* |
| RFK530 | *pid-2(tm1614); mjSi22[Pmex-5::mCherry::his-58::21UR-1_as::tbb-2(3'UTR)] (RNAe) I* |
| RFK586 | *pid-2(tm1614); mjSi22[Pmex-5::mCherry::his-58::21UR-1_as::tbb-2(3'UTR)] (RNAe) I* |
| SX2078 | *mjSi22[Pmex-5::mCherry::his-58::21UR-1_as::tbb-2(3'UTR)] I* |
| RFK416 | *mjSi22[Pmex-5::mCherry::his-58::21UR-1_as::tbb-2(3'UTR)] I; hrde-1(tm1200) III* |
| RFK587 | *pid-2(tm1614) I; prg-1(n4357) I* |
| RFK709 | *pid-2(xf23) I* |
| NL3643 | *unc-22(st136::Tc1) IV* |
| RFK611 | *wago-1(tm1414) I; wago-2(tm2686) I; ppw-2(tm1120) I; unc-22::Tc1(st136::Tc1) IV* |
| RFK610 | *pid-2(xf23) I; unc-22::Tc1(st136::Tc1) IV* |
| RFK612 | *pid-2(xf23) I; prg-1(n4357) I; unc22::Tc1 (st136::Tc1) IV* |
| RFK614 | *pid-2(xf23) I; hrde-1(tm1200) III; unc22::Tc1(st136::Tc1) IV* |
| RFK616 | *pid-2(tm1614) I; unc22::Tc1(st136::Tc1) IV* |
| RFK617 | *pid-2(tm1614) I; prg-1(n4357) I; unc22::Tc1(st136::Tc1) IV* |
| RFK619 | *pid-2(tm1614) I; hrde-1(tm1200) III; unc22::Tc1(st136::Tc1) IV* |
| SX523 | *prg-1(n4357) I* |
| RFK315 | *mjSi22[Pmex-5::mCherry::his-58::21UR-1_as::tbb-2(3'UTR)] I; pid-1(xf35) II; otIs45[unc-119::GFP] V* |
| RFK804 | *pid-2(xf23) I; mjSi22[Pmex-5::mCherry::his-58::21UR-1_as::tbb-2(3'UTR)] I; pid-1(xf35) II; otIs45[unc-119::GFP] V* |
| RFK182 | *pid-1(xf35) II* |
| RFK774 | *pid-2(xf23) I; pid-1(xf35) II* |
| | *xfls83[Pgl-3(5'UTR)::3xFLAG::pid-2::tbb-2(3'UTR); cb-unc119(+)] II* |
| | *xfls98[Pgl-3(5'UTR)::3xFLAG::pid-2::tbb-2(3'UTR); cb-unc119(+)] V* |
| RFK504 | *pid-2(xf23) I; xfls83[Pgl-3(5'UTR)::3xFLAG::pid-2::tbb-2(3'UTR); cb-unc119(+)] II; mjls144[Pmex-5::egfp::his-58::21UR-1_as::tbb-2(3'UTR)] II* |
| RFK505 | *pid-2(xf23) I; xfls98[Pgl-3(5'UTR)::3xFLAG::pid-2::tbb-2(3'UTR); cb-unc119(+)] V; mjls144[Pmex-5::egfp::his-58::21UR-1_as::tbb-2(3'UTR)] II* |
| RFK654 | *xfIs144[pid-2(5'UTR)::eGFP::pid-2::pid-2(3'UTR); cb-unc119(+)] II* |
| RFK655 | *xfIs145[pid-2(5'UTR)::pid-2::eGFP::pid-2(3'UTR); cb-unc119(+)] II* |
| RFK656 | *xfls146[pid-2(5'UTR)::3xFLAG::pid-2::pid-2(3'UTR); cb-unc119(+)] II* |
| RFK693 | *pid-2(xf23) I; xfls144[pid-2(5'UTR)::eGFP::pid-2::pid-2(3'UTR); cb-unc119(+)] II* |
| RFK694 | *pid-2(xf23) I; xfIs145[pid-2(5'UTR)::pid-2::eGFP::pid-2(3'UTR); cb-unc119(+)] II* |

| | |
|---|---|
| RFK695 | *pid-2(xf23) I; xfls146[pid-2(5'UTR)::3xFLAG::pid-2::pid-2(3'UTR); cb-unc119(+)] II* |
| RFK853 | *mjSi22[Pmex-5::mCherry::his-58::21UR-1_as::tbb-2(3'UTR)] I pid-2(xf23) I; xfls144[pid-2(5'UTR)::eGFP::pid-2::pid-2(3'UTR); cb-unc119(+)] II* |
| RFK854 | *mjSi22[Pmex-5::mCherry::his-58::21UR-1_as::tbb-2(3'UTR)] I pid-2(xf23) I; xfIs145[pid-2(5'UTR)::pid-2::eGFP::pid-2(3'UTR); cb-unc119(+)] II* |
| RFK855 | *mjSi22[Pmex-5::mCherry::his-58::21UR-1_as::tbb-2(3'UTR)] I pid-2(xf23) I; xfls146[pid-2(5'UTR)::3xFLAG::pid-2::pid-2(3'UTR); cb-unc119(+)] II* |
| RFK184 | *mjSi22[Pmex-5::mCherry::his-58::21UR-1_as::tbb-2(3'UTR)] I; pid-1(xf35) II* |
| RFK422 | *mjSi22[Pmex-5::mCherry::his-58::21UR-1_as::tbb-2(3'UTR)] I (RNAe); pid-1(xf35) II* |
| RFK764 | *mjSi22[Pmex-5::mCherry::his-58::21UR-1_as::tbb-2(3'UTR)] I; pid-1(xf35) II* |
| RFK765 | *mjSi22[Pmex-5::mCherry::his-58::21UR-1_as::tbb-2(3'UTR)] I; pid-1(xf35) II* |
| RFK766 | *mjSi22[Pmex-5::mCherry::his-58::21UR-1_as::tbb-2(3'UTR)] I; pid-1(xf35) II* |
| RFK767 | *mjSi22[Pmex-5::mCherry::his-58::21UR-1_as::tbb-2(3'UTR)] I (RNAe); pid-1(xf35) II* |
| RFK768 | *mjSi22[Pmex-5::mCherry::his-58::21UR-1_as::tbb-2(3'UTR)] I (RNAe); pid-1(xf35) II* |
| RFK769 | *mjSi22[Pmex-5::mCherry::his-58::21UR-1_as::tbb-2(3'UTR)] I (RNAe); pid-1(xf35) II* |
| RFK771 | *mjSi22[Pmex-5::mCherry::his-58::21UR-1_as::tbb-2(3'UTR)] I; pid-1(xf35) II; hrde-1(tm1200) III* |
| RFK857 | *pid-4(xf184) I* |
| RFK858 | *pid-4(xf185) I* |
| RFK859 | *pid-5(xf181) V* |
| RFK860 | *pid-5(xf182) V* |
| RFK847 | *pid-4(xf186[pid-4::3xMyc]) I* |
| RFK879 | *pid-5(xf192[pid-5::2xHA]) V* |
| RFK932 | *pid-4(xf204[pid-4::d10]) I* |
| RFK988 | *pid-4(xf206[pid-4::mTagRFP-T]) I* |
| RFK972 | *pid-5(xf221[pid-5::d10]) V* |
| RFK1025 | *pid-5(xf226[pid-5::mTagRFP-T]) V* |
| RFK982 | *pid-4(xf184) I; mjSi22[Pmex-5::mCherry::his-58::21UR-1_as::tbb-2(3'UTR)] I* |
| RFK983 | *pid-4(xf184) I; mjSi22[Pmex-5::mCherry::his-58::21UR-1_as::tbb-2(3'UTR)] (RNAe) I* |
| RFK984 | *mjSi22[Pmex-5::mCherry::his-58::21UR-1_as::tbb-2(3'UTR)] I; pid-5(xf181) V* |
| RFK985 | *mjSi22[Pmex-5::mCherry::his-58::21UR-1_as::tbb-2(3'UTR)] I; pid-5(xf181) (RNAe) V* |
| RFK986 | *pid-4(xf184) I; pid-5(xf181) V* |
| RFK987 | *pid-4(xf184) I; mjSi22[Pmex-5::mCherry::his-58::21UR-1_as::tbb-2(3'UTR)] I; pid-5(xf181) V* |
| RFK979 | *hrde-1(tm1200) III* |
| RFK1085 | *xfIs145[pid-2(5'UTR)::pid-2::eGFP::pid-2(3'UTR); cb-unc119(+)] II; pgl-1(xf233[pgl-1::mTagRFP-T]) IV* |
| RFK1194 | *deps-1(ax2063[deps-1::GFP]) I; pid-4(xf206[pid-4::mTagRFP-T]) I* |
| RFK1195 | *deps-1(ax2063[deps-1::GFP]) I; pid-5(xf226[pid-5::mTagRFP-T]) V* |
| RFK1119 | *pid-4(xf206[pid-4::mTagRFP-T]) I; znfx-1(gg544[3xFLAG::GFP::znfx-1]) II* |
| RFK1118 | *znfx-1(gg544[3xFLAG::GFP::znfx-1]) II; pid-5(xf226[pid-5::mTagRFP-T]) V* |
| RFK1186 | *pid-4(xf206[pid-4::mTagRFP-T]) I; xfIs145[pid-2(5'UTR)::pid-2::eGFP::pid-2(3'UTR); cb-unc119(+)] II* |
| RFK1078 | *xfIs145[pid-2(5'UTR)::pid-2::eGFP::pid-2(3'UTR); cb-unc119(+)] II; pid-5(xf226[pid-5::mTagRFP-T]) V* |
| RFK1184 | *znfx-1(gg544[3xFLAG::GFP::znfx-1]) II; pgl-1(xf233[pgl-1::mTagRFP-T]) IV* |

| | |
|---|---|
| RFK1185 | *pid-2(xf23) I; znfx-1(gg544[3xFLAG::GFP::znfx-1]) II; pgl-1(xf233[pgl-1::mTagRFP-T]) IV* |
| RFK1216 | *pid-4(xf184) I; znfx-1(gg544[3xFLAG::GFP::znfx-1]) II; pgl-1(xf233[pgl-1::mTagRFP-T]) IV; pid-5(xf181) V* |
| RFK1217 | *pid-4(xf184) I; znfx-1(gg544[3xFLAG::GFP::znfx-1]) II; pgl-1(xf233[pgl-1::mTagRFP-T]) IV* |
| RFK1218 | *znfx-1(gg544[3xFLAG::GFP::znfx-1]) II; pgl-1(xf233[pgl-1::mTagRFP-T]) IV; pid-5(xf181) V* |

**Microscopy**

20-25 worms have been picked to a drop of M9 (80 µl) on a slide, washed and then fixed with acetone (2 x 80 µl). After acetone has evaporated, worms have been washed 2 x 10 minutes with 80 µl of PBS-Triton X100 0,1%. After removing the excess of PBS-Triton X100 0,1%, the worms have been mounted on a coverslip with Fluoroshield™ with DAPI (5 µl) (Art. No. F6057, Sigma).

Alternatively, for live imaging, 20-25 worms have been picked to a drop of M9 (80 µl) on a slide, washed and then 2 µl of 1 M $NaN_3$ have been added to paralyze the worms. After removing the excess of M9, a slide prepared with 2% agarose (in water) has been placed on top of the coverslip and worms have been imaged directly.

Images have been acquired either at a Leica DM6000B microscope (objective HC PL FLUOTAR 20x 0.5 dry, Art. No. 11506503, Leica) or at a Leica TCS SP5 STED CW confocal microscope (objective HCX PL APO 'CS 63x / 1.2 water UV, Art. No. 11506280, Leica). Images have then been processed with Leica LAS software and ImageJ. Images representing the expression of PID-2, PID-4, PID-5, and P and Z granules markers have been processed with the Huygens Remote Manager v3.6 and deconvoluted (Huygens Deconvolution, SVI).

For scoring the 21U sensor as active or silenced, we have used a Leica M165FC widefield microscope. The 21U sensor has been scored as: active, if the fluorescence was easily visible with a lower magnification (Plan APO 1.0x, Art. No. 10450028; Leica); faint, if the fluorescence was only visible with a higher magnification (Plan APO 5.0x/0.50 LWD, Art. No. 10447243; Leica); silenced, if no fluorescence was visible. The worms have been later used also for live imaging with a Leica DM6000B microscope as described above.

Colocalization analysis

In order to perform colocalization analysis of the fluorescently tagged proteins PID-2, PID-4, PID-5 and of the P and Z granules markers, we used the DiAna plugin of (Fiji Is Just) ImageJ (Gilles et al., 2017; Schindelin et al., 2012). We used the deconvoluted images (Huygens Deconvolution, SVI), which consist of a single z-stack, and analysed the two fluorescence channels of interest at the time. First, we cropped an area containing 1-8 nuclei within the pachytene zone of the gonad arm, adding the area to the ROI manager, to ensure cropping of the same area in both channels being analysed. We then used DiAna_Segment to apply a mask to the images, considering all objects with size from 1 to 2000 pixels and then adjusted the threshold to ensure a faithful segmentation of the images. After segmentation, we performed the analysis using DiAna_Analyse and measured the surface area ($\mu m^2$), the distance between the two signals (µm) and the number of objects present in the cropped area. We then represented the distance between centres as a measure of colocalization. For each couple of fluorescent proteins, we analysed 4-10 images of gonads from individual animals.

**Small RNA sequencing**

Small RNA enrichment

In order to enrich for small RNAs, we used the *mir*Vana™ kit (Art. No. AM1561, Invitrogen™). 400 µl of *mir*Vana™ Lysis/Binding buffer and 48 µl of *mir*Vana™ Homogenate Additive have been added to the total RNA (80 µl). The mix has been incubated at RT for 5 minutes to denature RNA, then 1/3 of volume of 100% ethanol has been added and after mixing, samples have been spun down at 2500 x g for 4 minutes at RT to pellet large RNAs (>200 nt). The supernatant has been transferred to a new Eppendorf tube and RNA has been precipitated at -80 °C for 1 h with isopropanol (1:1). Samples have been centrifuged at maximum speed for at least 10 minutes at 4 °C to pellet small RNAs. The pellet has then been washed twice with 75% ethanol and spun down at maximum speed for 5 minutes at 4 °C. Pellet has been dried and resuspended in 16 µl of nuclease-free water. RNA quality has been checked at Nanodrop and on agarose gel and further processed for library preparation and deep sequencing.

Small RNA classification and quantification

Gene annotation was retrieved from Ensembl (release-38) and merged with transposon coordinates, retrieved from Wormbase (PRJNA13758.WS264), creating a custom annotation used for the analysis. Mapped reads were categorized in small RNA classes as follows: 21U RNAs are 21 nt long sequences mapping sense to annotated 21U RNA loci; 22G RNAs are 20-23 nt long and map antisense to protein-coding/pseudogenes/lincRNA/transposons; 26G RNAs, are those which are 26 nt long, and map antisense to annotated protein-coding/pseudogenes/lincRNA; and miRNAs are 20-24 nt long mapping sense to annotated miRNA loci. Read filtering was done with a python script (https://github.com/adomingues/filterReads/blob/master/filterReads/filterSmallRNAclasses.py) based on pysam v0.8.1 / htslib (Li et al., 2009), in combination with Bedtools intersect. For miRNA a stricter intersection was required (intersectBed -f 1.0). Reads belonging to each class were then counted for each library (total levels). For particular analysis the number of small RNAs in certain sub-classes was summarized based on previously published lists: ALG-3/-4 (Almeida et al., 2019a); ERGO-1 (Almeida et al., 2019a); CSR-1 (Conine et al., 2013); NRDE-3 (Zhou et al., 2014); mutators (Phillips et al., 2014); and WAGO-1 (Gu et al., 2009). The genomic locations of 22G were then intersected with that of the genes, and counted for each library.

22G RNAs coverage on 21U sensor

For targeting of the 21U sensor by 22G RNAs, we considered only sequences that were 22 nt long and mapping unambiguously to the 21U sensor sequence. Coverage was calculated with Bedtools v2.25.0 (genomeCoverageBed -ibam -d -strand "-") (Quinlan & Hall, 2010). Visualization was created with the R/Bioconductor package ggbio (Yin et al., 2012).

Secondary and tertiary 22G RNAs on 21U sensor

Tertiary populations were defined as the 22G RNAs mapping antisense to the mCherry coding sequence, within the 21U sensor (Sapetschnig et al., 2015). To define the secondary 22G (Sapetschnig et al., 2015), those surrounding the *21ur-1* recognition site, the spread from this site was visually estimated on a genome browser (IGV) to be +/- 200 bp. The reads mapping to these positions were counted and normalized to non-structural reads.

Coverage of local 22G RNAs on endogenous 21U RNA target sites

Following the analysis of (Lee et al., 2012), 21U RNA targets were identified by mapping the annotated 21U RNA sequences (WBCel235) to the genome with bowtie v1.2.1.1 (-n 1 -l 8 -e 300 -k 1000 –best), allowing for one mismatch in the 8 nucleotide seed and reporting up to 1000 valid alignments. The alignments were then filtered allowing for one T to G change in the seed, and two additional mismatches and one T to G change in the remaining alignment (see also Bagijn et al., 2012). These putative 21U RNA target coordinates were then intersected with the genomic location of WAGO-1 associated genes, to define a confident set of 21U RNA binding sites. Finally the 5’ 21U RNA mapping site was shifted by 10 nt (bedtools shift -s 10), extended in both directions by 50 bp (slopBed -l 50 -r 50 -s), and finally the 22G RNA coverage was calculated for these regions (coverageBed -d -s), and normalized by the number of non-structural reads.

Differential small RNA targeting

22Gs mapping to annotated features in the custom GTF were counted with htseq-count v0.9.0 (htseq-count -s reverse -f bam -m intersection-nonempty). Differential expression comparisons were performed with DESeq2 v.1.18.1 (Love et al., 2014). For the selection of genes differentially targeted (mRNA), a cut-off of at least a two-fold-change difference between conditions and an adjusted p-value less than 0.05 was applied.

Overlap with siRNA pathways

To determine if genes whose expression is affected in the *pid-2* mutants are shared with other siRNA pathways, we collected list of genes previously identified as being targeted by the following pathways: CSR-1 (Conine et al., 2013); NRDE-3 (Zhou et al., 2014); mutators (Phillips et al., 2014); WAGO-1 (Gu et al., 2009); ERGO-1 and ALG-3/-4 (Almeida et al., 2019a). ALG-3/-4 targets do not include those shared with ERGO-1 targets. See also Almeida at al., 2019a. Gene overlap significance was determined with the R/Bioconductor package GeneOverlap, and p-values calculated with Fisher’s exact test.

22G siRNA coverage

For the metagene analysis with 22G coverage over genes, we broadly followed the analysis described in (Ishidate et al., 2018). Briefly, 22G coverage, normalized to mapped non-structural reads, was plotted along the gene bodies with DeepTools computeMatrix scale-regions —metagene —transcriptID gene —transcript_id_designator gene_id —missingDataAsZero -b 0 -a 0 —regionBodyLength 2000 —binSize 20 —averageTypeBins mean. The resulting coverage matrix was used to produce the final plots in R. Coverage in each bin is normalized to the total 22G coverage of the gene. Genes with total coverage less than 5 were removed.

21U mCherry sensor sequence

2246-2733: mex-5; 2734-3640: mCherry; 3641-4012: his-58; 4040-4061: 21U; 4062-4393: tbb-2_UTR

ctttgagccaatttatccaagtccttgtaaaaagtattcgaaaattgttaacggataaaatgtttattataatatcaaaaacaatttgtcagttgacc
actttttgatatatttgacagaaaccgggatgaattggctcaaagttagggcgcccttctattacagggtttctgataacaaaccggttattaactc
ccaacaagggatgatttcaattcatcatgctcaaattgacccaaattaagttacatgacaaattcatcgcccttttcaactcttcttggctcatcatc
tgttattctgttcctattctgtctgcaccctataccccttttgcatactctctcgtcattcctttcttggatagtgtcttcttctccccagctctgctacttct
atgacttgcgcccgctgcttttccgctcgttcctctcctctctgacgtcttcttctgcttcttttctccttcctttttcccttctcaatcttcctctttttccca
tttctgtcaatcattcgaagaagaagaagaagaagaccctcattcatttcattttttttttctgtcggtgtgtgctgctcggttaagagtgagctctcta
ttccacgtcttcttctttttcttcttcgattcgaatcaatcactccaccaaaacgcattcgtttttgggattcaccccgcggttcgcaaTaggtttctttt
tcaaattattagcgttataaatagaaaaatggtggagtttcaataaaaatgataatttacaaagtgtattttgattacatgtactcaaaaaggttga
aatttttcataccagttttccggaaatccatctgatatcattatcgtatttttctttttaaaaatgttttcaaaaaaaaacaaaatatagctggttattt
ggcaccctctaattaccattttcctgtcacaccacactcttttccttcttccacttctttccgttttcagccgcttccaaccaaaccgatatgaaagccg
agcaacaacaatcgattccacccggctcggcgaccttcccgtcgcaggtgagactcagaaaactagagaaaccgctcaactaactcttgatatcc
gatttcattcttttctcttttctttttgttgaacttttccatatttccagatgccacggccaccaccaagcaccgaacaaggaatcacaacggaatcgg
agcttgcgaagaaagctcaaatcactccgaacgacgttttagcacttccgggaatcactcaaggtatcttgctccctttggatttcagtttaaaaca
taaatttaggattcttatgctccccatctgcgaacatctataacatcgagttcaccaagttccaaatccgtgatctggacactgagcaagtgcttttc
gagatcgccaaaccggagaacgatcaggagaatgatgagtcgccacaggagtcggcaagatacgtgcgttatagatttgctccaaactttttga
aactcaaaacggtcggagcaactgtggaattcaaagtaggagacatcccaatccatcatttccgaatgatcgaacgtcacttcttcaaagatcgc

cttctgaagtgttttgactttgaattcggattctgtattccgaattcacgaaacaactgtgaacatatctatgagttccctcaactctctcaacaactc
agtgagtcattattctaaaaagtacaattcaaaagactaatctcttttcagtggacgacatgatcaacaatccaaacgagactcgttctgacagc
ttctatttcgtcgataacaaactcgtcatgcacaacaaagccgactactcatatgatgcataaatatttaatacaaaaatgttctggataattattct
gtcgaatagaaaaaaaactccaaatgtgattaaattccaataattcctgtctagtttgttccttccccttccccttctcatgttcaatgcattcctaag
cttttcagttccccccttgtttctatatttttcgctgtcctgtcacactcgctaaaaacactaatcacacggaaatctgttttcaataaaaactccaac
tttaactcattttcaatttcaactgaaagatttttcattagagaatgtctagaactagGCCCGGGCTACGTAATACGACTCACTTAAG
GCCTTGACTAGAGGGTACCAGAGCTCACCTAGGcaggaacagctatgaccatgattacgccaagctatcaactttGTATAGA
AAAGTTGAAATATCAGTTTTTAAAAAATTAAACCATAAAACAAATAATATAACCCAATTTTTACATCAAACCACA
AGAAAAAAATACATTTGGGCCCACGGATAAAGAAATTAAAAAAATACATTTTTTAAAGGCGCACCGAATTAAA
ATTCATTTGGGTCTTACCGCGTATACCGTACTCCGTTTGTTTGATCATTTTTGTCAGCGCTGGCGGTTGTTTTTC
ATTTCATTTCTGCTTCAAAGACGTTTTCTCGAATAATTTTTCGTTTATTCTCTTTTTTAAAATTAATTTCTAGCCGT
AAATGTTATAAATTCACCCATTTAACGCAAATTTCATGGTAATCTCATGGAAAAATGCAGTTTCTTTGTTAAAGA
AAGCTTAAATAGCAAAAATTCCCCGACTTTCCCCAAAATCCTGCTCGATTTTCCGTTTTCTCATTGTATTCTCTCT
TAATTAATTTTATCGATAATCAATTGAATGTTTCAGACAGAGAATGGTCTCAAAGGGTGAAGAAGATAACATG
GCAATTATTAAAGAGTTTATGCGTTTCAAGGTGCATATGGAGGGATCTGTCAATGGGCATGAGTTTGAAATTG
AAGGTGAAGGAGAAGGCCGACCATATGAGGGAACACAAACCGCAAAACTAAAGgtaagtttaaacatatatatactaa
ctaaccctgattatttaaattttcagGTAACTAAAGGCGGACCATTACCATTCGCCTGGGACATCCTCTCTCCACAGTTCA
TGTATGGAAGTAAAGCTTATGTTAAACATCCGGCAGATATACCAGATTATTTGAAACTTTCATTCCCGGAGGGT
TTTAAGTGGGAACGCGTAATGAATTTTGAAGACGGAGGAGTTGTTACAGTGACGCAAGACTCAAGgtaagtttaa
acagttcggtactaactaaccatacatatttaaattttcagCCTCCAAGATGGAGAATTTATTTATAAAGTCAAACTTCGAGGA
ACGAATTTCCCCTCGGATGGACCTGTTATGCAGAAGAAGACTATGGGATGGGAAGCTTCAAGTGAAAGAATG
TACCCTGAAGACGGTGCTCTTAAGGGAGAGATTAAACAACGTCTTAAATTGAAAGATGGAGGACATTACGATG
CTGAGgtaagtttaaacatgattttactaactaactaatctgatttaaattttcagGTGAAGACAACTTACAAAGCCAAAAAACCA
GTTCAGCTGCCAGGAGCGTACAATGTTAATATTAAACTGGATATCACCTCCCACAACGAGGATTACACTATCGT
TGAGCAATATGAAAGAGCTGAAGGGCGGCACTCGACAGGTGGCATGGATGAATTGTATAAGGGAGGTGGAG
GTGGAGCTTCAAGTTTGTACAAAAAAGCAGGCTCGATGCCACCAAAGCCATCTGCCAAGGGAGCCAAGAAGG
CCGCCAAGACCGTCGTTGCCAAGCCAAAGGACGGAAAGAAGAGACGTCATGCCCGCAAGGAATCGTACTCCG
TCTACATCTACCGTGTTCTCAAGCAAGTTCACCCAGACACCGGAGTCTCCTCCAAGGCCATGTCTATCATGAACT
CCTTCGTCAACGATGTATTCGAACGCATCGCTTCGGAAGCTTCCCGTCTTGCTCATTACAACAAACGCTCAACG
ATCTCATCCCGCGAAATTCAAACCGCTGTCCGTTTGATTCTCCCAGGAGAACTTGCCAAGCACGCCGTGTCTGA
GGGAACCAAGGCCGTCACCAAGTACACTTCCAGCAAGTAAACCCAGCTTTCTTGTACAAAGTGGtcaagcacggtt
aacgtacgtaccaGATAAATGCAAAATCCTTTCAAGCATTCCCTTCTTCTCTATCACTCTTCTTTCTTTTTGTCAAAAA
ATTCTCTCGCTAATTTATTTGCTTTTTTAATGTTATTATTTTATGACTTTTTATAGTCACTGAAAAGTTTGCATCTG
AGTGAAGTGAATGCTATCAAAATGTGATTCTGTCTGATGTACTTTCACAATCTCTCTTCAATTCCATTTTGAAGT
GCTTTAAACCCGAAAGGTTGAGAAAAATGCGAGCGCTCAAATATTTGTATTGTGTTCGTTGAGTGACCCAACA
AAAAGAGGAAACTTTATTGTGCCGCCAAGAAAAAAGTCTCACAACTTTATTATACAAAGTTGataattcactggccgt
cgttttacaCTCGAGACGTACGGTGCGCGCGATGCATTCGAAGATCTGCCCACTAGTGAGTCGTATTA

**Production of PID-2 protein for antibody generation**

<u>Cloning</u>

In order to produce a polyclonal antibody against PID-2, we cloned the full-length coding sequence (CDS) of *pid-2* (UniProtID Q9N3P1) into a vector for recombinant protein overexpression in *E. coli*.

The cDNA, obtained from wild type worms with the ProtoScript® First Strand cDNA Synthesis Kit (Art. No. E6300S, New England Biolabs), has been used as template for the amplification of the CDS of *pid-2* using Q5® Hot Start High-Fidelity DNA Polymerase (Art. No. M0493, New England Biolabs). PCR primers were designed to contain restriction sites (forward: NcoI ggccatgggcatgacagttattatagcgtcacact; reverse (without stop codon): XhoI ggctcgagaaatggg cactcgctgaat) for subsequent cloning into the pET-28a(+) expression vector (Art. No. 69864-3, Merck), which confers bacterial resistance to Kanamycin and adds a 6x histidine tag for affinity purification, either at the N- or at the C-terminus of the protein of interest. The PCR product was purified from agarose gel (QIAquick Gel Extraction Kit, Art. No. 28706, QIAgen). The vector and the PCR product (insert) have been digested with the restriction enzymes NcoI (Art. No. R3193, New England Biolabs) and XhoI (Art. No. R0146, New England Biolabs) for 2 hours at 37 °C. The insert has been additionally dephosphorylated using Antarctic Phosphatase (Art. No. M0289, New England Biolabs) for 15 minutes at 37 °C to avoid self-ligation. Ligation was then performed for 30 minutes at room temperature using a molar ratio 3:1 = vector:insert using T4 DNA Ligase (Art. No. M0202, New England Biolabs). The ligation reaction was then transformed in Subcloning Efficiency™ DH5α™ Competent Cells (Art. No. 18265017, Invitrogen™) and plated for selection on LB agar plates with Kanamycin (30 µg/ml). The plasmid was isolated from the bacterial culture (PureLink™ HiPure Plasmid Miniprep Kit, Art. No. K210011, Invitrogen™), checked by enzymatic digestion and sequencing.

<u>Protein expression and purification</u>

For protein expression, the pET-28a(+)-PID-2 construct has been transformed into Rosetta™ (DE3) Competent Cells (Art. No. 70954-3, Merck) and positive clones have been selected on LB agar plates with Kanamycin (30 µg/ml). A single colony has been inoculated into 20 ml of LB media supplied with 100 µg/ml Kanamycin and 35 µg/ml Chloramphenicol as pre-culture and grown overnight at 37 °C. The pre-culture has then been inoculated in 1 l LB media supplied with 100 µg/ml Kanamycin and 35 µg/ml Chloramphenicol and grown for 3 hours at 37 °C, until exponential phase (OD600 = 0,6). Protein expression has been induced by adding 0,5 mM IPTG (Art. No. V3953, Promega) and incubated overnight at 18 °C. The bacterial culture has been harvested by spinning down at 4000 x g for 15 minutes at 4 °C. Cells pellet has been collected and washed with HEPES buffer to remove excess of growth media. The bacteria were spun down again at 4500 x g for 20 minutes at 4 °C and the pellet was frozen at -20 °C. In order to purify PID-2 protein tagged with 6xHis-tag from inclusion bodies, bacterial pellet was thawn on ice and resuspended in 30 ml of lysis buffer (500 mM NaCl; 100 mM Tris HCl pH=8.5). The pellet has been homogenized, sonicated 3 x 3 minutes (Branson Sonifier 450; output 4-5; duty cycle 2-3) and the cell lysate has been centrifuged at 19000 x g for 25 minutes at 4 °C. The pellet containing PID-2 has then been resuspended in 30 ml of denaturing buffer (500 mM NaCl; 100 mM Tris HCl pH=8.5; 8 M urea) to solubilize the inclusion bodies and centrifuged again at 19000 x g for 25 minutes at 4 °C to remove cells debris. The supernatant has then been loaded on a batch column containing 1 ml of Ni-NTA Agarose slurry (Art. No. 30210, QIAgen), previously equilibrated with the same buffer (500 mM NaCl; 100 mM Tris HCl pH=8.5; 8 M urea). After binding of the protein, the Ni-NTA Agarose beads have been washed with 10 ml of buffer and then eluted with 10 ml of elution buffer (500 mM NaCl; 100 mM TrisHCl pH=8.5; 4 M Urea; 250 mM imidazole). Eluate was stored at -80 °C.

<u>Antibody production</u>

After checking the purity of the protein by SDS-PAGE, the eluted protein PID-2::6xHis has been concentrated to a final concentration of 1,8 mg/ml and sent to Eurogentec for antibody production (two rabbits; 28-day Speedy protocol). We then received the serum from two rabbits (823 and 824) and used 823 for all the experiments (1:100 for immunoprecipitation).

**Transgenic lines generation using the miniMos system**

We generated *pid-2* transgenic lines using the miniMos system, as previously described (Frøkjær-Jensen et al., 2014). The injection mix contains plasmids encoding for the co-injection markers (10 ng/µl pGH8; 2,5 ng/µl pCFJ90; 5 ng/µl pCFJ104), for the transposase (50 ng/µl pCFJ601) and for the desired transgene embedded in a modified Mos element (10 ng/µl pRK1012). We injected the mix in the strain HT1593, which carries an *unc-119(ed3) III* mutation, and kept the worms at 25 °C until starvation. We first screened for mCherry expressing worms, indicative of a successful injection, and later on, for wild-type moving worms that have no extrachromosomal array (no mCherry expression), indicating that the *unc-119(ed3) III* mutation has been rescued by the wild-type allele encoded by the template plasmid pRK1012. We have isolated seven independent insertions of the desired transgene. Worms were then lysed and genotyped to confirm the insertion of the transgene. Afterwards, we mapped the insertion of the transgenes, using an inverse PCR approach, as previously described (Frøkjær-Jensen et al., 2014). We managed to map only two insertions, *xfIs83* and *xfIs98*. The insertion of the transgene *xfIs83* was mapped to chromosome II, within the last intron of *mpz-1*, whereas the transgene *xfIs98* was mapped to chromosome V, within the fourth intron of the Y32B12B.4 gene.

**Transgenic lines generation using the MosSCI system**

In order to avoid potential variability effects on the expression of the *pid-2* transgenes generated with the miniMos system, due to their insertion locus, we generated *pid-2* transgenic lines, using the MosSCI system and targeted the locus *ttTi5605* on LGII, as previously described (Frøkjær-Jensen et al., 2008). The injection mix contains plasmids encoding for the co-injection markers (10 ng/µl pGH8; 2,5 ng/µl pCFJ90; 5 ng/µl pCFJ104), for the transposase (50 ng/µl pCFJ601) and for the desired transgene (50 ng/µl pRK1036, pRK1037 or pRK1038). We injected the mix in the strain EG6699, which carries a Mos insertion on the locus *ttTi5605* on LGII, and kept the worms at 25 °C until starvation. We first screened for mCherry expressing worms, indicative of a successful injection, and later on, for wild-type moving worms that have no extrachromosomal array (no mCherry expression). From the templates pRK1036, pRK1037 and pRK1038, we have isolated *xfIs144* [eGFP::PID-2], *xfIs145* [PID-2::eGFP] and *xfIs146* [3xFLAG::PID-2], respectively. Worms were then lysed and genotyped to confirm the insertion of the transgene.

**Generation of mutant and endogenously tagged lines using CRISPR/Cas9 technology**

Generation of mutant lines

Wild type worms have been injected with an injection mix containing 50 ng/µl pJW1259 (encoding for *Peft-3::cas9::tbb-2 3'UTR*, a gift from Jordan Ward (Ward, 2015)), co-injection markers (10 ng/µl pGH8; 5 ng/µl pCFJ104; 2,5 ng/µl pCFJ90) and 30 ng/µl of each of the plasmids encoding for the sgRNAs, specifically pRK1054, pRK1056, pRK1057, pRK1059 to target the *Y45G5AM.2* locus and pRK1047, pRK1050, pRK1052, pRK1053 to target the *W03G9.2* locus. After injections, worms have been kept at 20 °C and F1 offspring expressing the co-injection markers have been singled out. After the F1 offspring have laid embryos, they have been picked to 5 µl of single worm lysis buffer and the lysate has been used as PCR template to screen for mutant alleles. We isolated two deletion alleles of *Y45G5AM.2/pid-5* (*xf181* and *xf182*) and two deletion alleles of *W03G9.2/pid-4* (*xf184* and *xf185*). Each allele has been sequenced to pinpoint the exact deletion at nucleotide resolution. The mutant strains have been outcrossed two times against wild type N2 strain to remove any potential off-targets effect of Cas9 and used for further experiments.

Generation of endogenously tagged lines

In order to introduce an epitope tag at the endogenous loci of *W03G9.2/pid-4* and *Y45G5AM.2/pid-5*, we used the co-conversion approach as previously described (Arribere et al., 2014). Wild type worms have been injected with an injection mix containing 50 ng/µl pJS164 (Cas9 + sgRNA *dpy-10*); 750 nM ssODN SJ665 (repair oligo for *dpy-10*(*cn64*)); 50 ng/µl pRK1053; 750 nM ssODN SJ969 (repair oligo for W03G9.2::3xMyc) to introduce a 3xMyc epitope tag at the endogenous *W03G9.2/pid-4* locus (*xf186*). Wild type worms have been injected with an injection mix containing 50 ng/µl pJS164; 500 nM ssODN SJ665; 50 ng/µl pRK1060; 1000 nM ssODN SJ964 (repair oligo for Y45G5AM.2::2xHA) to introduce a 2xHA epitope tag at the endogenous *Y45G5AM.2/pid-5* locus (*xf192*). After injections, worms have been kept at 20 °C and F1 offspring with a roller phenotype (*rol-6*) have been singled out. After the F1 offspring have laid embryos, they have been picked to 5 µl of single worm lysis buffer and the lysate has been used as PCR template to screen for edited alleles. Each allele has been sequenced to ensure that the insertion of the epitope is in frame. The tagged strains have been outcrossed two times against wild type N2 strain to remove any potential off-targets effect of Cas9 and used for further experiments.

To introduce a fluorescent protein at the endogenous locus, we have first used a *unc-58* co-conversion approach (Arribere et al., 2014) to introduce at the *W03G9.2/pid-4* and at the *Y45G5AM.2/pid-5* loci a sequence of 20 nucleotides of *dpy-10* gene that serves as efficient protospacer sequence for subsequent edits (*xf204* and *xf221*, respectively), as previously described (Mouridi et al., 2017). We then used the generated strains (RFK932 and RFK972) as reference for the injection of a mix containing 50 ng/µl pJS164 (Cas9 + sgRNA *dpy-10*); 1000 nM SJ665 (repair oligo for *dpy-10*(*cn64*)); 300 ng/µl SJP010, a PCR product that was amplified from plasmid pDD286 (a gift from Bob Goldstein, Addgene plasmid # 70684) and used as a donor for the insertion of the mTagRFP-T sequence at the endogenous *W03G9.2/pid-4* and *Y45G5AM.2/pid-5* loci (Paix et al., 2014). After injections, worms have been kept at 20 °C and F1 offspring with a roller phenotype have been singled out. After the F1 offspring have laid embryos, they have been picked to 5 µl of single worm lysis buffer and the lysate has been used as PCR template to screen for edited alleles. We isolated the *xf206* and *xf226* alleles, which have been sequenced to ensure that the insertion of the mTagRFP-T sequence is in frame. The tagged strains have been outcrossed two times against wild type N2 strain to remove any potential off-targets effect of Cas9 and used for further experiments.

<u>List of primers and plasmids generated</u>

| Plasmid | Oligo | Sequence |
|---|---|---|
| | p46169_RV | aaacatttagatttgcaatt |
| pRK1047 | W03G9.2_sgRNA2 | gcatgaacacgcataacgcgagttttagagctagaaatagc |
| pRK1050 | W03G9.2_sgRNA5 | gcaaagtacgcgaaagagggttttagagctagaaatagc |
| pRK1052 | W03G9.2_sgRNA7 | gcttcatatagctgccgctcggttttagagctagaaatagc |
| pRK1053 | W03G9.2_sgRNA8 | gaatatcaagcactagttggcgttttagagctagaaatagc |
| pRK1054 | Y45G5AM.2_sgRNA1 | ggctgtcatcagcgctcgtgttttagagctagaaatagc |
| pRK1056 | Y45G5AM.2_sgRNA3 | gtttcgcggttcgaagctacggttttagagctagaaatagc |
| pRK1057 | Y45G5AM.2_sgRNA4 | ggattggaatcaacgtgaggttttagagctagaaatagc |
| pRK1059 | Y45G5AM.2_sgRNA6 | gaatgagatcgatcgaagccggttttagagctagaaatagc |
| pRK1060 | Y45G5AM.2_sgRNA7 | caattaaaaatgctctagatgtttaagagctatgctggaaac |

<u>List of ssODN repair templates</u>

SJ665
cacttgaacttcaatacggcaagatgagaatgactggaaaccgtaccgcatgcggtgcctatggtagcggagcttcacatggcttcagaccaac agcctat

SJ969
gatgaaaaataattaaagcttgaatatcaagcactacaagtcttcctcgctgatcaacttctgctcgaggtcctcctcggagatgagcttttgctca agatcctcttcagaaataagtttttgttcacctccacctccggatccgttggccggcttcatatagctgccgctcgcgggaa

SJ964
cgaaaataacttaaaaacaattaaaaatgctctaggcatagtctggaacgtcatatgggtaagcgtaatctgggacatcgtatggataacctcca cctccggatccgatcggctggcatgcattgaagagccagtcatattc

**Protein extraction and immunoprecipitation using GFP-Trap®**

Synchronized worms have been grown until adulthood, then washed with cold M9 buffer, collected in a final volume of 200 µl water and fast frozen on dry ice. 350 µl of 2X Lysis Buffer (20 mM Tris pH=7.5; 300 mM NaCl; 1 mM EDTA; 1% NP40; cOmplete™ Mini EDTA-free Protease Inhibitor Cocktail, Art. No. 11836170001, Roche) have been added to each sample. Worms have then been sonicated using Bioruptor Plus (30 seconds ON – 30 seconds OFF, 10 cycles, high) and lysates were spun down at maximum speed for 10 min at 4 °C. The cleared protein extracts (500 µl) have been transferred to a new tube and 30 µl of GFP-Trap®_M (Art. No. gtm-20, ChromoTek), previously equilibrated with Dilution/Wash Buffer (3 x 5 minutes) (10 mM Tris pH=7.5; 150 mM NaCl; 0,5 mM EDTA; cOmplete™ Mini EDTA-free Protease Inhibitor Cocktail), have been added to each sample and the immunoprecipitation was performed for 3 hours at 4 °C. Then, beads have been washed 3 x 5 minutes with 500 µl of Dilution/Wash Buffer. After the last wash, the beads have been resuspended in 25 µl of NuPAGE® LDS Sample Buffer 1x (Art. No. NP0007, Life technologies) with 100 mM DTT and boiled at 95 °C for 10 minutes.

**Protein extraction and immunoprecipitation using Dynabeads™ Protein G**

Synchronized worms have been grown until adulthood, then washed with cold M9 buffer, collected in a final volume of 200 µl water and fast frozen on dry ice. 350 µl of 2X Lysis Buffer (50 mM Tris HCl pH=7.5; 300 mM NaCl; 3 mM $MgCl_2$; 2 mM DTT; 0,2% Triton-X100; cOmplete™ Mini EDTA-free Protease Inhibitor Cocktail, Art. No. 11836170001, Roche) have been added to each sample. Worms have then been sonicated using Bioruptor Plus (30 seconds ON – 30 seconds OFF, 10 cycles, high) and lysates were spun down spun down at maximum speed for 10 min at 4 °C. The cleared protein extracts (500 µl) have been transferred to a new tube and 2 µg of antibody (αHA clone HA-7, Art. No. H3663, Sigma; αMYC 9B11, Art. No. 2276, Cell Signalling Technology) or 1:100 of serum (823 αPID-2) have been added to each sample. The immunoprecipitation was performed for 2 hours at 4 °C. Then, 30 µl of Dynabeads™ Protein G (Art.No. 10004D, Invitrogen™), previously equilibrated with Wash Buffer (3 x 5 minutes) (25 mM Tris HCl pH=7.5; 150 mM NaCl; 1,5 mM $MgCl_2$; 1 mM DTT; cOmplete™ Mini EDTA-free Protease Inhibitor Cocktail), have been added and incubate for 1 additional hour at 4 °C. Then, beads have been washed 3 x 5 minutes with 500 µl of Wash Buffer. After the last wash, the beads have been resuspended in 25 µl of NuPAGE® LDS Sample Buffer 1x (Art. No. NP0007, Life technologies) with 100 mM DTT and boiled at 95 °C for 10 minutes.

**Mass Spectrometry**

Label-free quantitative mass spectrometry has been performed as described in (Almeida et al., 2018). After boiling (see above), the samples were separated on a 4–12% gradient Bis-Tris gel (NuPAGE Bis-Tris gels, 1.0 mm, 10 well; Art. No. NP0321; Life Technologies) in 1X MOPS (NuPAGE 20X MOPS SDS running buffer; Art. No. NP0001; Life Technologies) at 180 V for 10 min, afterwards processed by in-gel digest (Bluhm et al., 2019; Shevchenko et al., 2007) and desalted using a C18 StageTip (Rappsilber et al., 2007). The digested peptides were separated on a 25-cm reverse-phase capillary (75 µM inner diameter) packed with Reprosil C18 material (Dr. Maisch) with a 2 h gradient from 2 to 40% Buffer B (see Stage tip purification) with the EASY-nLC 1,000 system (Thermo Scientific). Measurement was done on a Q Exactive Plus mass spectrometer (Thermo Scientific) operated with a Top10 data-dependent MS/MS acquisition method per full scan (Bluhm et al., 2016). The measurements were processed with the MaxQuant software, version 1.5.2.8 (Cox & Mann, 2008) against the Wormbase *C. elegans* database (version of WS265) for quantitation and the Ensemble *E. coli* REL606 database (Version Oct 2018) to filter potential contaminations.

**Protein extraction and Western blot**

For each strain, 50 L4 worms were hand-picked, to ensure comparable protein concentration and the right larval stage, in 13 µl of M9 buffer 1X. Subsequently, 5 µl of NuPAGE™ LDS Sample Buffer 4X (Art.

No. NP0007, Invitrogen™) and 2 µl of DTT 1M have been added. Samples have been boiled at 95 °C for 10 minutes and then stored at -20 °C, until all samples have been collected.

Protein extracts have been loaded, together with a broad range protein ladder (Color Prestained Protein Standard, Broad Range; Art. No. # P7712; NEB), onto a 10% Bis-Tris gel (NuPAGE 10% Bis-Tris Protein Gels, 1.0 mm, 10 well; Art. No. NP0301BOX; Invitrogen™) and run with MOPS buffer 1X (NuPAGE™ MOPS SDS Running Buffer (20X); Art. No. NP0001; Invitrogen™), at 140 V at 4 °C. Afterwards, proteins have been transferred on a PVDF membrane (Immobilon-P Membrane, PVDF, 0.45 µm; Art. No. IPVH00010; Merck Millipore) for 90 min, at 100 V at 4 °C, with Transfer buffer 1X (NuPAGE™ Transfer Buffer (20X); Art. No. NP00061; Invitrogen™) plus 10% methanol (Methanol, anhydrous 99.8%; Art. No. 322415-2L; Sigma-Aldrich Chemie GmbH). The membrane has been incubated with blocking buffer (5% milk in PBS-Tween 0,1%) for 1 h at room temperature. The membrane has then been cut, according to the molecular weight of the proteins to be detected, and each part has been incubated with a dilution of the primary antibody in 0,5% milk in PBS-Tween 0,1% (1:5000 Monoclonal ANTI-FLAG® M2 antibody produced in mouse, Art. No. F3165, Sigma-Aldrich; 1:5000 RFP Rabbit anti-Tag, Polyclonal, Art. No. 10041338, Fisher Scientific GmbH; 1:2500 Monoclonal anti-α-Tubulin antibody produced in mouse, Art. No. T6074, Sigma-Aldrich) for 1 h at room temperature. Then, three washes of five minutes each have been done with PBS-Tween 0,1% at room temperature. The three parts of the membrane have then been incubated with a dilution of the secondary antibody (1:10000 Anti-mouse IgG, HRP-linked Antibody, Art. No. #7076, Cell Signaling Technology; 1:10000 Anti-rabbit IgG, HRP-linked Antibody, Art. No. #7074, Cell Signaling Technology) in PBS-Tween 0,1% for 1 h at room temperature. Afterwards, three washes of five minutes each have been done with PBS-Tween 0,1% at room temperature, and the blot has been developed using the Amersham ECL Select Western Blotting Detection Reagent (Art. No. RPN2235, GE Healthcare) and detected at the ChemiDoc XRS+ (Bio-Rad).

**Data availability**

Sequencing data is available at NCBI-SRA, BioProject ID PRJNA612883. Mass spectrometry data has been deposited at PRIDE, accession number PXD018402.

**SUPPLEMENTAL FIGURE LEGENDS.**

**Figure S1.**

**A)** Scheme of the single-copy MosSCI and miniMos transgenes expressing tagged PID-2. The MosSCI transgenes (*xfIs144, xfIs145, xfIs146*) are driven by the endogenous promoter and 3' UTR and were inserted into a known germline-expressing site on chromosome II, *ttTi5605*. The miniMos transgenes (*xfIs83, xfIs98*) instead are driven by a germline specific promoter (*pgl-3*) and 3'UTR (*tbb-2*) and have been randomly inserted in the genome. The *xfIs83* transgene was mapped to chromosome II, within the last intron of *mpz-1*, whereas the transgene *xfIs98* was mapped to chromosome V, within the fourth intron of the Y32B12B.4 gene.

**B)** Crossing strategy to assess the rescue of the *pid-2* mutation by the indicated *pid-2* transgenes.

**C)** Quantification of the expression of the 21U sensor (% of animals analysed) in the indicated mutant backgrounds as a measure of the rescue of the *pid-2* mutation by the different *pid-2* transgenes. Expression was quantified by scoring individuals (ON/OFF) using microscopy.

**D)** Representative images of the 21U sensor expression (left: mCherry signal; right: DIC) in the indicated genetic backgrounds. For *xfIs145*, both ON and OFF states are depicted. Gonads are outlined by a dashed line. Scale bar: 25 µm.

**E)** RT-qPCR of the 21U sensor in the indicated mutant backgrounds. *: strains that received the 21U sensor from a *mut-7* background (active). §: strains that received a silenced 21U sensor from a *prg-1* background (RNAe). Expression of *pmp-3* was used to normalize the data. Error bars reflect the standard deviation, calculated from three technical replicates. We note that the *pid-2* strain, that received the 21U sensor originally in an RNAe-state, has RNA levels of the 21U sensor comparable to a de-silenced sensor in a *pid-2* mutant background. Indeed, upon re-examination of this strain under the microscope, we detected expression of the 21U sensor, indicating that in this isolate RNAe had been lost (also see Figure S1F).

**F)** Quantification of the reactivation of the 21U sensor (indicated as % of plates containing animals with detectable expression), at either 20 °C or at 25 °C, in the indicated mutant backgrounds. From each of the indicated mutant backgrounds, 20 L2-L3 larvae were singled out and scored to ensure that the 21U sensor was still silenced (RNAe). Then, 10 plates were kept at 20°C and 10 plates were kept at 25°C and chunked regularly to avoid starvation. After 14 days, plates were scored by microscopy for re-activation of the 21U sensor. *pid-2(tm1614)*$_1$: RFK530; *pid-2(tm1614)*$_2$: RFK586.

**G)** 22G RNAs profile on the 21U sensor, schematically represented at the bottom, in the indicated mutant backgrounds. The top two panels refer to control strains, whereas the bottom two panels show profiles from strains isolated from an individual in which the 21U sensor was exposed to maternal 21U RNAs only. Both silenced as well as non-silenced strains were sequenced. In each plot, the average of three biological replicates is represented. The shading represents the standard deviation among the replicates.

**H)** Representative images of the 21U sensor expression (left: mCherry signal; right: DIC) in the indicated genetic backgrounds. The two top panels represent strains that have been exposed to maternal 21U RNAs only. In the strain depicted on top, the sensor became silenced, whereas in the strain depicted below it did not get silenced. Upon introduction of *hrde-1* mutation in the strain carrying a 21U sensor silenced upon exposure to maternal 21U RNAs only, the 21U sensor is reactivated, as shown at the bottom. Gonads are outlined by a dashed line. Scale bar: 25 µm.

**I)** Examples of *pid-1;pid-2* double mutant animals, isolated from a growing *pid-1*;*pid-2* double mutant population, showing feminization (upper panel) and pseudo-males (lower panel). The arrow indicates arrayed appearance of oocytes in the feminized animal (upper panel), while it indicates a male-like tail in the pseudo-male (lower panel). The latter also shows characteristics of hermaphrodites (two gonad arms, eggs in uterus and vulva). Scale bar: 100 µm.

**Figure S2.**

**A)** Representation of the total abundance of small RNA classes (21U, 22G, 26G RNAs and miRNAs) from small RNA sequencing data of the indicated genetic backgrounds. Each replicate is represented by a dot and the median is represented by a bar. P-values are calculated with a two-tailed unpaired t-test. RPM: reads per million. N2: wild type.

**B)** Dot plot for quantification of secondary (around the 21U RNA recognition site; upper panel) and tertiary (within the mCherry coding region; lower panel) 22G RNAs complementary to the 21U sensor in the indicated genetic backgrounds. Each replicate is represented by a dot and the median is represented by a bar. P-values are calculated with a two-tailed unpaired t-test. RPM: reads per million.

**C)** Violin plots representing the distribution of different sub-types of 22G RNAs as previously defined, as a group, in the indicated genetic backgrounds. The white boxes inside each of the violin plots represent the $75^{th}$ and $25^{th}$ percentile of the distribution, top and bottom respectively. The median of the distribution is represented by the line in each box. P-values are calculated with a two-sided unpaired Mann-Whitney/Wilcoxon rank-sum test, indicating the differences between *pid-2* mutants and either wild type or *prg-1* mutants as references. RPM: reads per million.

**D)** Profile of antisense 22G RNAs produced in a 100bp window around endogenous 21U RNA target sites (+50 bp; -50 bp) of WAGO-1 target genes, centred on position 10 of the 21U RNA sequence. Each line represents a specific genotype and the shading represents the standard deviation of the biological triplicates for the indicated genetic backgrounds.

**Figure S3.**

**A-D)** UpSet plots showing the overlap between genes up- and down-regulated in the *pid-2* mutants and known genes targeted from different small RNA pathways.

**Figure S4.**

**A, B)** Volcano plot representing the enrichment of proteins interacting with PID-2, as determined by immunoprecipitation of 3xFLAG::PID-2 protein, followed by quantitative label-free mass spectrometry. Results from two strains with independently generated miniMos transgenes (*xfIs83* in **A** and *xfIs98* in **B**) are shown. As control, FLAG IPs were performed on extracts from wild-type, non-transgenic animals. For each IP-MS experiment, quadruplicates were measured and analysed. The dashed line reflects a significance threshold of p-value < 0.05 at two-fold enrichment.

**C)** Multiple sequence alignment, using ESPript 3 (http://espript.ibcp.fr) (Robert & Gouet, 2014), of the predicted eTudor domains of PID-5 and PID-4 with eTudor domains of proteins from other organisms. If the conservation is >70%, the columns are framed in blue. The residues perfectly conserved are highlighted in red. The residues written in black are not conserved. Dots (.) indicate gaps in the protein sequence generated through the alignment. The residues characteristic of eTudor domains are indicated by the arrow (eTD). The aromatic residues forming the aromatic cage are highlighted in green, and the conserved acidic amino acid residue is highlighted in yellow. Ce: *Caenorhabditis elegans*; Dm: *Drosophila melanogaster*; Bm: *Bombyx mori*; Mm: *Mus musculus*; Hs: *Homo sapiens*.

**D)** Multiple sequence alignment of the X-Prolyl aminopeptidase domain of PID-5 (CePID-5) with its orthologs, and with APP-1 (CeAPP-1) and APP-1 orthologs from other nematodes, using ESPript 3 (http://espript.ibcp.fr) (Robert & Gouet, 2014). Conservation >70% is framed in blue. Perfectly conserved residues are highlighted in red. The residues in black are not conserved. Dots indicate gaps in the protein sequence generated through the alignment. The dashed line separates the protein according to the presence or absence of the catalytic residues, which are underlined. *C. brenneri*: CBN14695; *C. remanei*: CRE09268; *C. briggsae*: CBG01441; *S. ratti*: SRAE_2000061700; *P. pacificus*: Ppa-APS-3 PPA08577; *B. malayi*: Bm6170; *C. japonica*: Cjp-APP-1.

**E)** Genomic locus and predicted transcript (Wormbase) for *pid-4*. The alleles generated using CRISPR/Cas9 technology are indicated. *xf184*: -1570 + 8 nt; *xf185*: -1587 nt; *xf186*: 3xMyc; *xf206*: mTagRFP-T.

**F)** Genomic locus and predicted transcripts (Wormbase) for *pid-5*. The alleles generated using CRISPR/Cas9 technology are indicated. *xf181*: -5291 + 5 nt; *xf182*: -114 + 3 nt; *xf192*: 2xHA; *xf226*: mTagRFP-T.

**G, H)** Volcano plots representing the enrichment of proteins interacting with PID-2, as determined by immunoprecipitation of the endogenous PID-2 protein, followed by quantitative label-free mass spectrometry. IPs on protein extracts from *pid-2* mutant were compared to IPs on protein extracts from *pid-4* (**G**) and *pid-5* (**H**) mutants. For each IP-MS experiment, quadruplicates were measured and analysed. The dashed line reflects a significance threshold of p-value < 0.05 at two-fold enrichment.

**Figure S5.**

**A)** Plots representing 22G RNA reads mapping against the 21U sensor, schematically represented at the bottom, in *pid-4* and *pid-5* mutant backgrounds. The 21U sensor was crossed into the two mutant backgrounds in either a silenced state, from a *prg-1* mutant (RNAe), or in an active state, from a *mut-7* mutant (not indicated as RNAe). Both *mut-7* and *prg-1* were confirmed to be wild type in the strains that were isolated from these crosses. In each plot, the average of three biological replicates is represented and the shading represents the standard deviation among the replicates.

**B)** Representation of the total abundance of small RNA classes (21U, 22G, 26G RNAs and miRNAs) from small RNA sequencing of the indicated genetic backgrounds. Each replicate is represented by a dot and the median is represented by a bar. P-values are calculated with a two-tailed unpaired t-test. RPM: reads per million.

**C)** Violin plots representing the distribution of different sub-types of 22G RNAs as previously defined, as a group, in the indicated genetic backgrounds. The white boxes inside each of the violin plots represent the $75^{th}$ and $25^{th}$ percentile of the distribution, top and bottom respectively. The median of the distribution is represented by the line in each box. P-values are calculated with a two-sided unpaired Mann-Whitney/Wilcoxon rank-sum test, indicating the differences between *pid-2* mutants and either wild type or *prg-1* mutants as references. RPM: reads per million.

**Figure S6.**

**A-B)** Cumulative 22G coverage along the gene body of different sets of genes. Values represent 22G coverage normalized to the total coverage of the gene, for wild type (N2) and *pid-4* (**A**) or *pid-5* (**B**) mutants, of the previously defined 22G RNA target sub-types: ALG-3/-4 (Almeida et al., 2019a), CSR-1 (Conine et al., 2013), ERGO-1 (Almeida et al., 2019a), mutator (Phillips et al., 2014). The lines represent the average of biological replicates, whereas the shading represents the standard deviation of biological replicates. a.u.: arbitrary units.

**Figure S7.**

**A)** Boxplots representing the area of Z granules ($\mu m^2$) in wild type, *pid-2*, *pid-4/-5*, *pid-4* and *pid-5* mutant backgrounds. The area of each Z granule is represented by a dot and the median is represented by a bar. The $75^{th}$ and $25^{th}$ percentile are represented by the upper and lower lines, respectively. P-values were calculated using a t-test (two-tailed).

**B)** Box-plots representing the distance (µm) between P and Z granules, in wild type, *pid-2*, *pid-4;pid-5*, *pid-4* and *pid-5* mutant backgrounds. The distance between each pair of fluorescent proteins is represented by a dot and the median is represented by a bar. The $75^{th}$ and $25^{th}$ percentile are represented by the upper and lower lines, respectively. P-values were calculated using a t-test (two-tailed).

**C, D)** Tables listing the number of PID-4::mTagRFP-T (**D**) and PID-5::mTagRFP-T (**E**) foci and the number of nuclei used for quantification in a wild-type and in a *pid-2* mutant background, represented in **Figure 6D, E** and **7G, H**. For each cropped area, the number of nuclei and foci was counted, and statistical differences between wild type and *pid-2* mutants were determined with a one-tailed weighted Student's t-test, implemented in the R package "weights", which accounts for the different number of nuclei in each area,. P-values are indicated in the tables.

**E)** Western blot to detect the expression of 3xFLAG::GFP::ZNFX-1, using αFLAG antibody, and PGL-1::mTagRFP-T, using αRFP antibody, in the indicated strains. Per lane, 50 L4 larvae were used to make a lysate by boiling in NuPAGE® LDS sample buffer. αTubulin has been used as loading control.

**Table S1.** List of genes differentially targeted in *pid-2* mutants, compared to wild type, generated with DESeq2. To obtain the genes highlighted in blue and red in **Figure 3A**, filter for fold-changes in the column “log2 fold change (MLE): group pid2 mut vs WT” and for adjusted p-value in the column “BH adjusted p-values”.

**Table S2.** List of genes differentially targeted in *pid-4* mutants, compared to wild type, generated with DESeq2. To obtain the genes highlighted in blue and red in **Figure 5B**, filter for fold-changes in the column “log2 fold change (MLE): group pid4 mut vs WT” and for adjusted p-value in the column “BH adjusted p-values”.

**Table S3.** List of genes differentially targeted in *pid-5* mutants, compared to wild type, generated with DESeq2. To obtain the genes highlighted in blue and red in **Figure 5D**, filter for fold-changes in the column “log2 fold change (MLE): group pid5 mut vs WT” and for adjusted p-value in the column “BH adjusted p-values”.

Figure S1

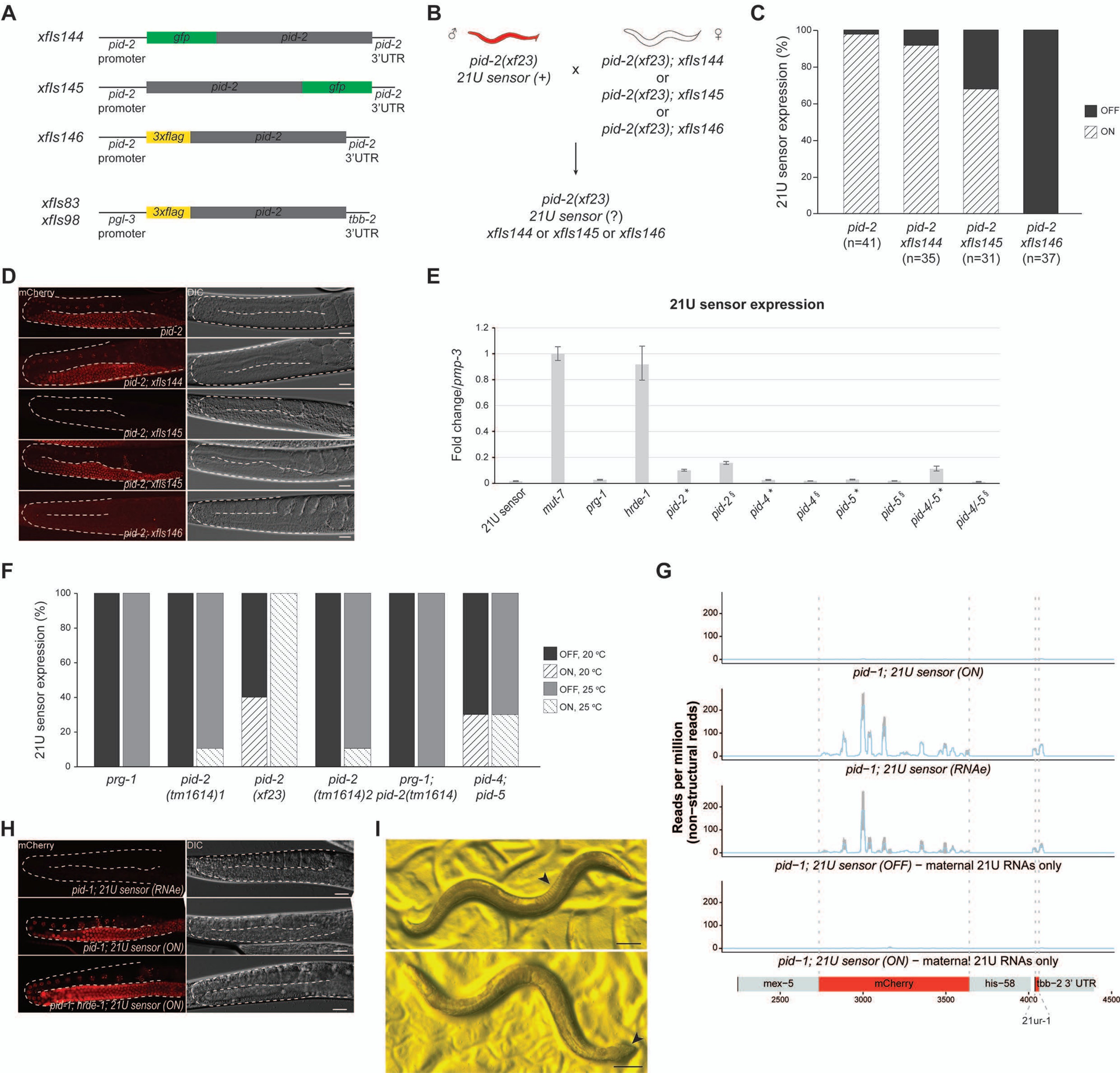

A
xfIs144
pid-2 promoter
gfp
pid-2
pid-2 3'UTR
xfIs145
xfIs146
3xflag
xfIs83
xfIs98
pgl-3 promoter
tbb-2 3'UTR
B
pid-2(xf23)
21U sensor (+)
x
pid-2(xf23); xfIs144
or
pid-2(xf23); xfIs145
or
pid-2(xf23); xfIs146
pid-2(xf23)
21U sensor (?)
xfIs144 or xfIs145 or xfIs146
C
21U sensor expression (%)
OFF
ON
pid-2 (n=41)
pid-2 xfIs144 (n=35)
pid-2 xfIs145 (n=31)
pid-2 xfIs146 (n=37)
D
mCherry
DIC
pid-2
pid-2; xfIs144
pid-2; xfIs145
pid-2; xfIs146
E
21U sensor expression
Fold change/pmp-3
21U sensor
mut-7
prg-1
hrde-1
pid-2 *
pid-2 §
pid-4 *
pid-4 §
pid-5 *
pid-5 §
pid-4/-5 *
pid-4/-5 §
F
21U sensor expression (%)
OFF, 20 ºC
ON, 20 ºC
OFF, 25 ºC
ON, 25 ºC
prg-1
pid-2 (tm1614)1
pid-2 (xf23)
pid-2 (tm1614)2
prg-1; pid-2(tm1614)
pid-4; pid-5
G
Reads per million (non-structural reads)
pid−1; 21U sensor (ON)
pid−1; 21U sensor (RNAe)
pid−1; 21U sensor (OFF) − maternal 21U RNAs only
pid−1; 21U sensor (ON) − maternal 21U RNAs only
mex−5
mCherry
his−58
tbb−2 3' UTR
21ur-1
H
mCherry
DIC
pid-1; 21U sensor (RNAe)
pid-1; 21U sensor (ON)
pid-1; hrde-1; 21U sensor (ON)
I

Figure S2

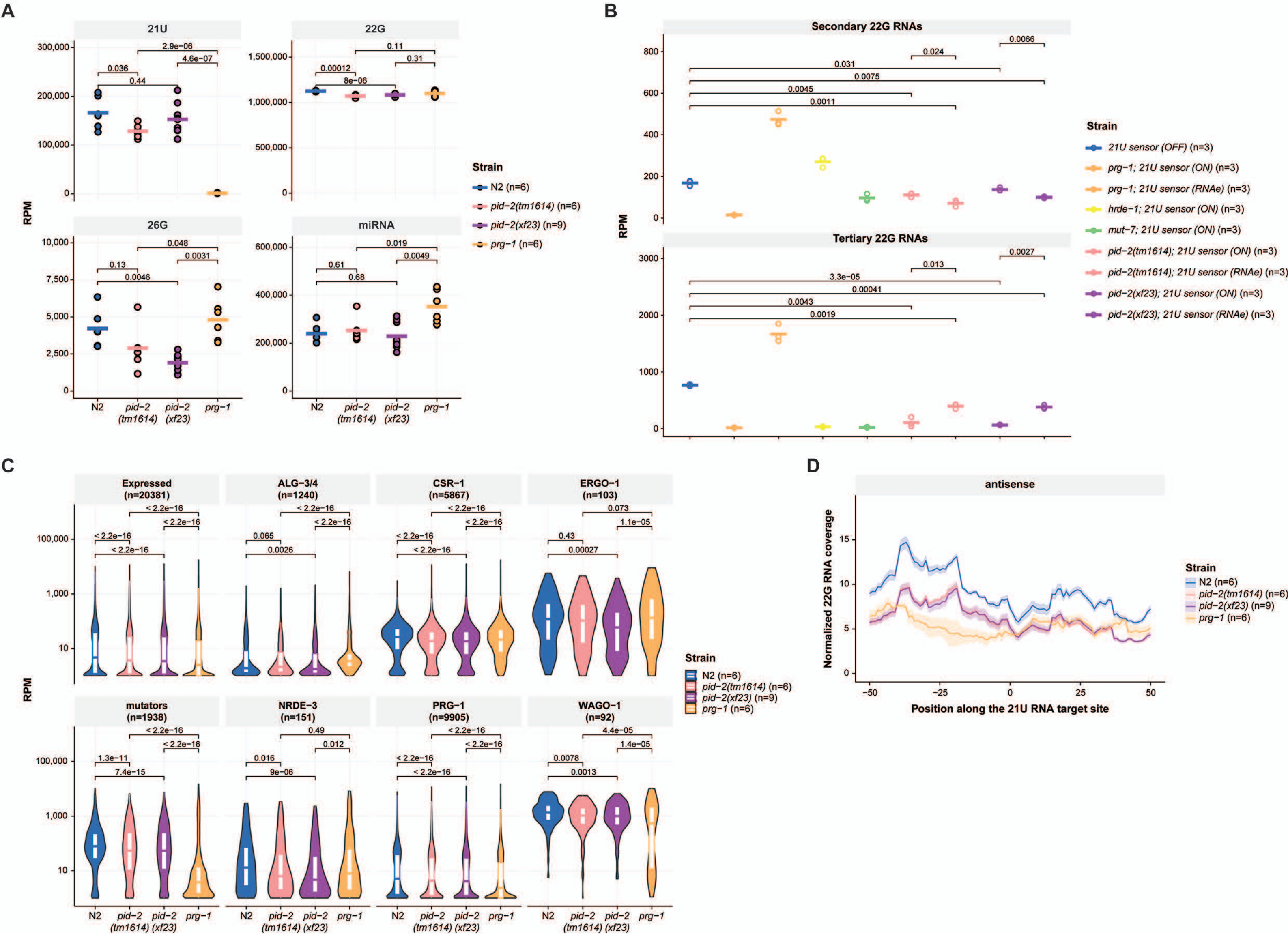

A
21U
22G
26G
miRNA
RPM
N2
pid−2 (tm1614)
pid−2 (xf23)
prg−1
Strain
N2 (n=6)
pid−2(tm1614) (n=6)
pid−2(xf23) (n=9)
prg−1 (n=6)
B
Secondary 22G RNAs
Tertiary 22G RNAs
RPM
Strain
21U sensor (OFF) (n=3)
prg−1; 21U sensor (ON) (n=3)
prg−1; 21U sensor (RNAe) (n=3)
hrde−1; 21U sensor (ON) (n=3)
mut−7; 21U sensor (ON) (n=3)
pid−2(tm1614); 21U sensor (ON) (n=3)
pid−2(tm1614); 21U sensor (RNAe) (n=3)
pid−2(xf23); 21U sensor (ON) (n=3)
pid−2(xf23); 21U sensor (RNAe) (n=3)
C
Expressed (n=20381)
ALG−3/4 (n=1240)
CSR−1 (n=5867)
ERGO−1 (n=103)
mutators (n=1938)
NRDE−3 (n=151)
PRG−1 (n=9905)
WAGO−1 (n=92)
RPM
Strain
N2 (n=6)
pid−2(tm1614) (n=6)
pid−2(xf23) (n=9)
prg−1 (n=6)
D
antisense
Normalized 22G RNA coverage
Position along the 21U RNA target site
Strain
N2 (n=6)
pid−2(tm1614) (n=6)
pid−2(xf23) (n=9)
prg−1 (n=6)

Figure S3

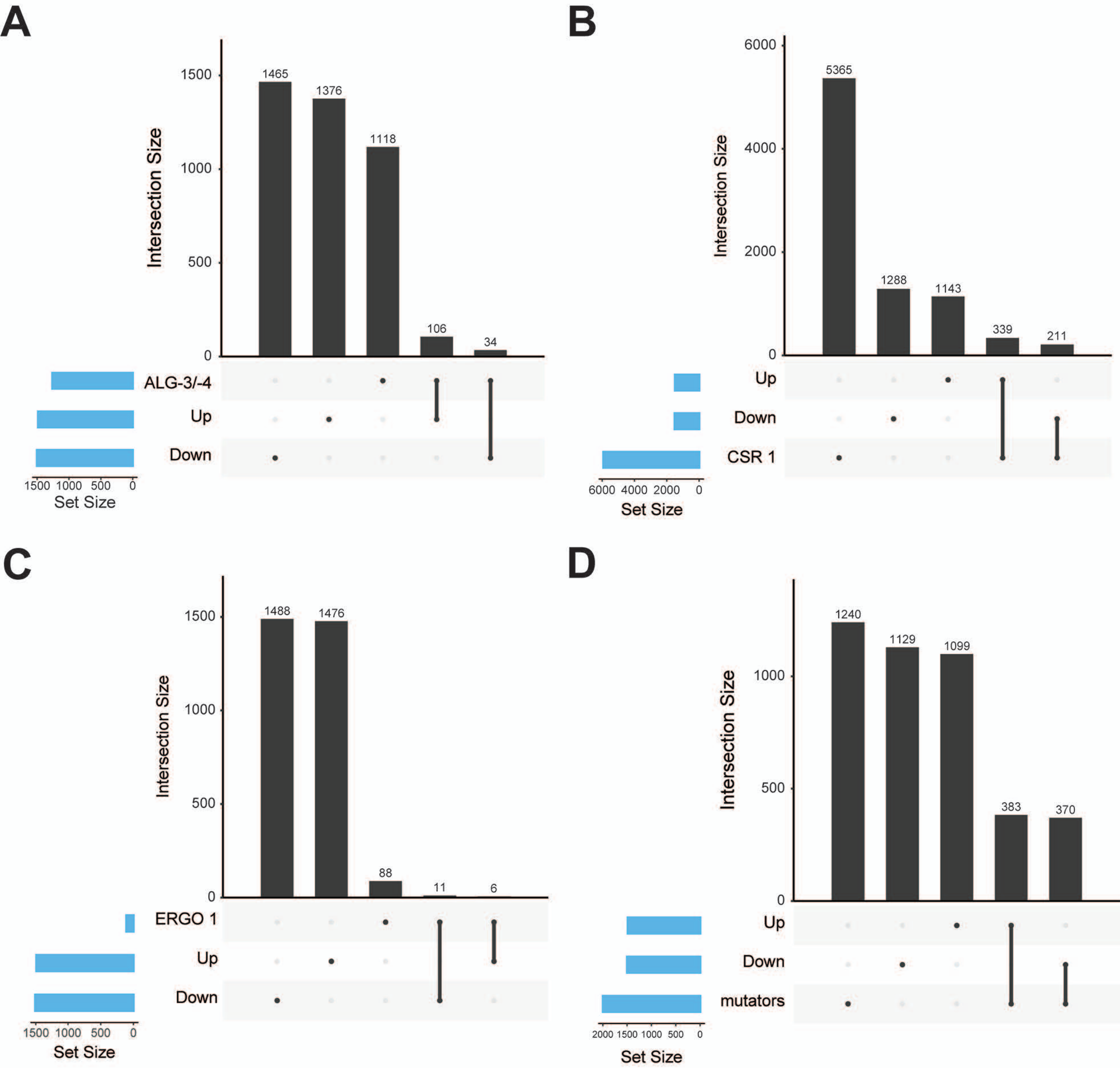
A
1465
1376
1118
106
34
Intersection Size
1500
1000
500
0
ALG-3/-4
Up
Down
1500 1000 500 0
Set Size
B
5365
1288
1143
339
211
Intersection Size
6000
4000
2000
0
Up
Down
CSR 1
6000 4000 2000 0
Set Size
C
1488
1476
88
11
6
Intersection Size
1500
1000
500
0
ERGO 1
Up
Down
1500 1000 500 0
Set Size
D
1240
1129
1099
383
370
Intersection Size
1000
500
0
Up
Down
mutators
2000 1500 1000 500 0
Set Size

Figure S4

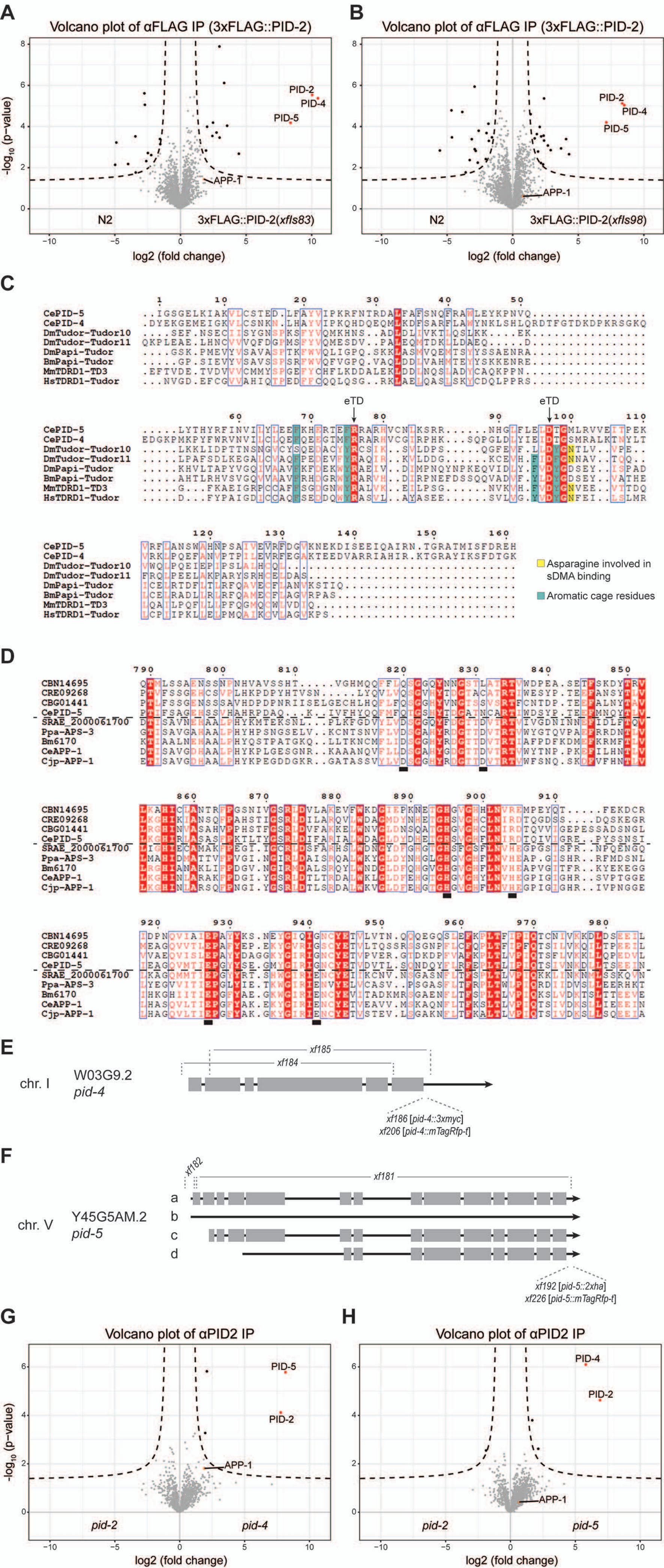
A
Volcano plot of αFLAG IP (3xFLAG::PID-2)
-log10 (p-value)
PID-2
PID-4
PID-5
APP-1
N2
3xFLAG::PID-2(xfIs83)
log2 (fold change)
B
Volcano plot of αFLAG IP (3xFLAG::PID-2)
PID-2
PID-4
PID-5
APP-1
N2
3xFLAG::PID-2(xfIs98)
log2 (fold change)
C
CePID-5
CePID-4
DmTudor-Tudor10
DmTudor-Tudor11
DmPapi-Tudor
BmPapi-Tudor
MmTDRD1-TD3
HsTDRD1-Tudor
eTD
eTD
Asparagine involved in sDMA binding
Aromatic cage residues
D
CBN14695
CRE09268
CBG01441
CePID-5
SRAE_2000061700
Ppa-APS-3
Bm6170
CeAPP-1
Cjp-APP-1
E
xf185
xf184
chr. I
W03G9.2
pid-4
xf186 [pid-4::3xmyc]
xf206 [pid-4::mTagRfp-t]
F
xf182
xf181
a
b
c
d
chr. V
Y45G5AM.2
pid-5
xf192 [pid-5::2xha]
xf226 [pid-5::mTagRfp-t]
G
Volcano plot of αPID2 IP
-log10 (p-value)
PID-5
PID-2
APP-1
pid-2
pid-4
log2 (fold change)
H
Volcano plot of αPID2 IP
PID-4
PID-2
APP-1
pid-2
pid-5
log2 (fold change)

Figure S5

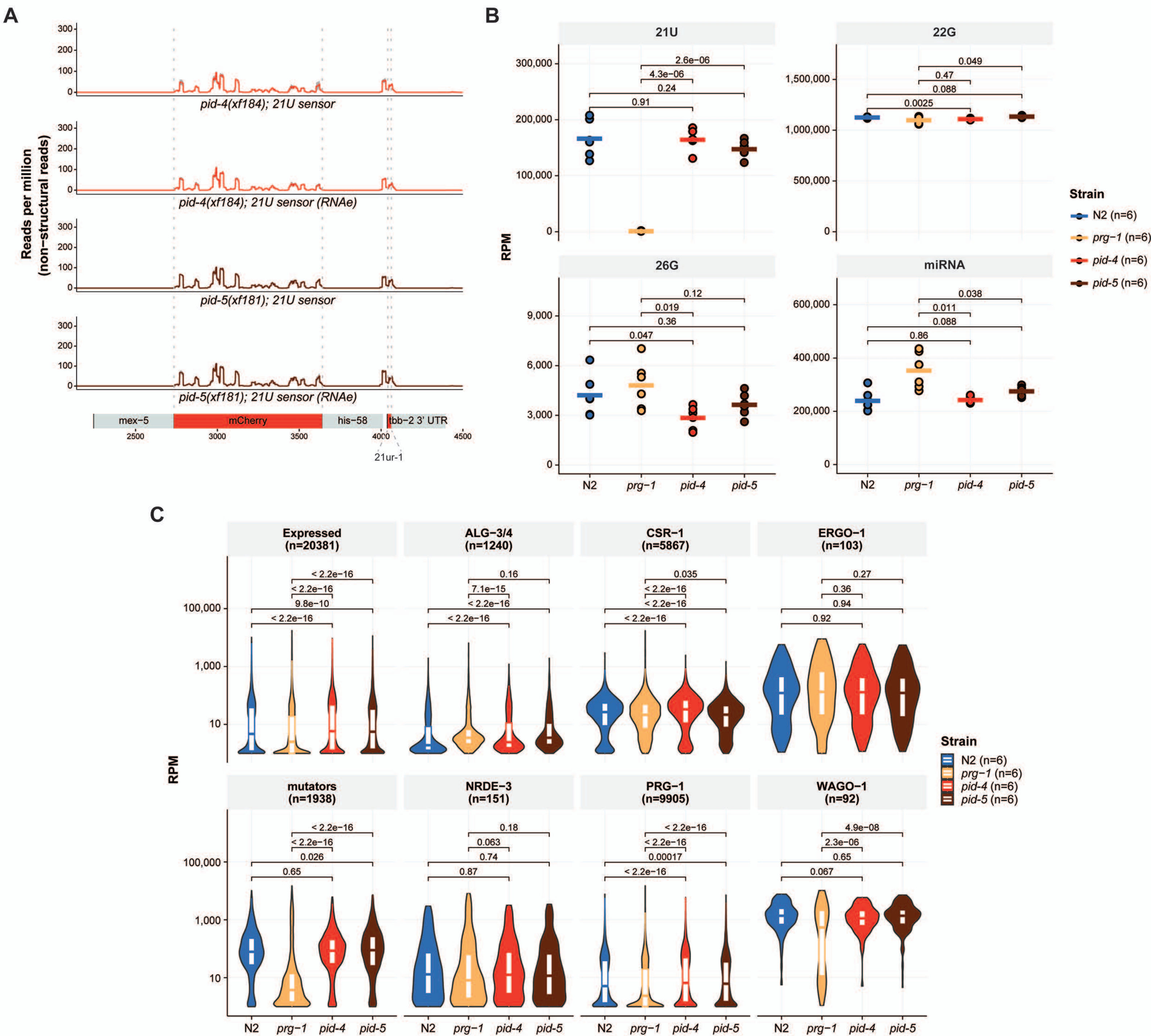
A
Reads per million (non-structural reads)
pid-4(xf184); 21U sensor
pid-4(xf184); 21U sensor (RNAe)
pid-5(xf181); 21U sensor
pid-5(xf181); 21U sensor (RNAe)
mex-5
mCherry
his-58
tbb-2 3' UTR
21ur-1
B
21U
22G
26G
miRNA
RPM
N2
prg-1
pid-4
pid-5
Strain
N2 (n=6)
prg-1 (n=6)
pid-4 (n=6)
pid-5 (n=6)
C
Expressed (n=20381)
ALG-3/4 (n=1240)
CSR-1 (n=5867)
ERGO-1 (n=103)
mutators (n=1938)
NRDE-3 (n=151)
PRG-1 (n=9905)
WAGO-1 (n=92)
RPM
Strain
N2 (n=6)
prg-1 (n=6)
pid-4 (n=6)
pid-5 (n=6)

Figure S6

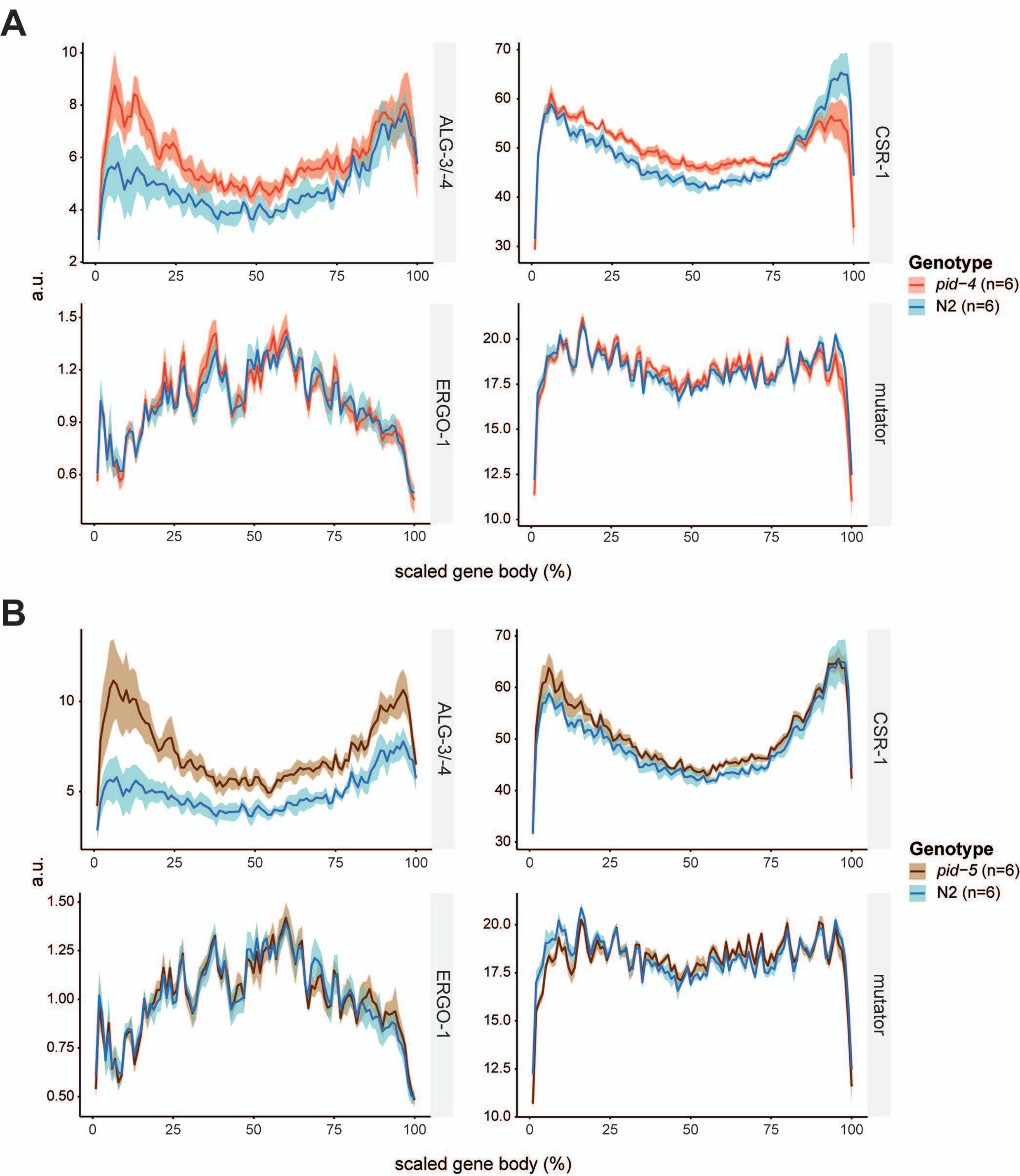
A
a.u.
ALG-3/-4
CSR-1
ERGO-1
mutator
scaled gene body (%)
Genotype
pid−4 (n=6)
N2 (n=6)
B
a.u.
ALG-3/-4
CSR-1
ERGO-1
mutator
scaled gene body (%)
Genotype
pid−5 (n=6)
N2 (n=6)

Figure S7

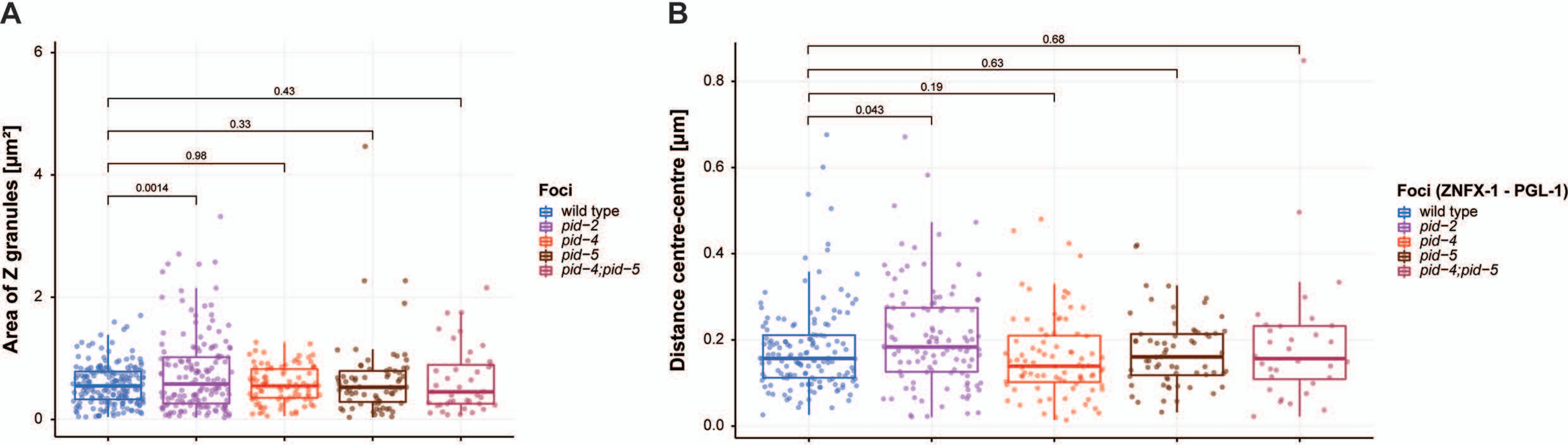


**C**

| **PID-4::mTagRFP-T** (p-value = 0.014) | | |
|---|---|---|
| | **wild type** | ***pid-2*** |
| **Total nuclei** | 23 | 26 |
| **Total foci** | 169 | 122 |
| **Average number foci / nucleus** | 7.3 | 4.7 |

**D**

| **PID-5::mTagRFP-T** (p-value = 0.36) | | |
|---|---|---|
| | **wild type** | ***pid-2*** |
| **Total nuclei** | 24 | 25 |
| **Total foci** | 80 | 69 |
| **Average number foci / nucleus** | 3.3 | 2.8 |

**E**

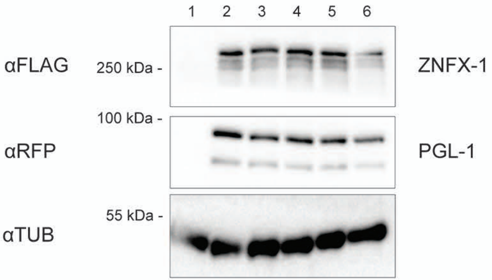


1: N2
2: *znfx-1(gg544[3xflag::gfp::znfx-1]); pgl-1(xf233[pgl-1::mTagRfp-t])*
3: *znfx-1(gg544[3xflag::gfp::znfx-1]); pgl-1(xf233[pgl-1::mTagRfp-t]); pid-2(xf23)*
4: *znfx-1(gg544[3xflag::gfp::znfx-1]); pgl-1(xf233[pgl-1::mTagRfp-t]); pid-4(xf184); pid-5(xf181)*
5: *znfx-1(gg544[3xflag::gfp::znfx-1]); pgl-1(xf233[pgl-1::mTagRfp-t]); pid-4(xf184)*
6: *znfx-1(gg544[3xflag::gfp::znfx-1]); pgl-1(xf233[pgl-1::mTagRfp-t]); pid-5(xf181)*
